# Phage terminase recognition by the bacterial immune sensors Avs2 and Upx

**DOI:** 10.64898/2026.04.30.721860

**Authors:** Simone A. Evans, Collin Chiu, Max E. Wilkinson, David Li, Mahamaya Biswal, Jonathan Strecker, Tino Pleiner, Feng Zhang, Alex Gao

## Abstract

Prokaryotes employ diverse defense strategies to detect and halt the progression of phage infection. Multiple defense systems sense phage proteins through direct binding, including antiviral STAND NTPases (Avs), which oligomerize upon target recognition to induce programmed cell death. The widespread Avs2 family was previously shown to detect the large terminase subunit of tailed phages, but the mechanism of terminase sensing was unknown. Here, we determine the structural basis of terminase recognition by Avs2 from *Escherichia coli* (EcAvs2). A cryo-EM structure at 2.3 Å resolution reveals that EcAvs2 forms a flat, C4-symmetric tetramer in which each protomer is bound to a single terminase monomer. Terminase recognition is mediated by a large, shape complementary binding pocket in the EcAvs2 sensor domain, including specific contacts with an unexpected ATP molecule at the interface of EcAvs2 and terminase. Furthermore, we demonstrate that the defense protein Upx also recognizes diverse phage terminases, despite lacking sequence and structural homology to Avs. AlphaFold 3 models indicate that Upx binds an unfolded state of the core terminase ATPase domain, mediated by β-augmentation. These findings highlight the distinct modes of terminase recognition across structurally diverse defense proteins.

## Introduction

Innate immune systems across all domains of life detect molecular signatures of pathogen infection^1,2^. In prokaryotes, multiple lines of defense intercept every stage of the phage life cycle, from initial entry to the expression of viral proteins or the disruption of host translation. A recurring strategy across these defenses is the direct detection of phage proteins by dedicated sensor modules^3–15^. However, how structurally distinct sensors achieve detection of diverse phage protein homologs while maintaining self−non-self specificity is not fully understood.

Several defense systems target the highly conserved large terminase subunit (hereafter, terminase), the central component of the DNA packaging motor of tailed phages. Consisting of ATPase and nuclease domains, the terminase cleaves concatemeric phage DNA into genome-sized fragments and translocates them into capsids^16^. To date, three antiviral STAND (Avs) proteins, Avs1, Avs2, and Avs3, have been identified as terminase sensors. Avs1−3 differ from defense systems such as TerI^15^ and Rip1 (ref.^13^) that sense the distinct, non-enzymatic small terminase subunit. These Avs proteins share a conserved tripartite domain architecture: a C-terminal sensor that binds the phage protein, a central ATPase that facilitates oligomerization, and an N-terminal effector that subsequently induces cell death to halt phage propagation.

Previously, we demonstrated that *Salmonella enterica* Avs3 (SeAvs3) tetramerizes upon terminase binding, activating its nuclease domain to degrade dsDNA. SeAvs3 recognizes immutable features of the terminase, including active site residues and its bound ATP ligand, contributing to its ability to detect a broad range of terminases with minimal sequence similarity^3^. While Avs1 shares strong predicted structural similarity with Avs3, Avs2 is predicted to be structurally distinct, and how it recognizes terminases remains unknown. Avs2 is also the most widespread and abundant of these systems, with extensive horizontal gene transfer and effector domain diversity^3^.

In addition, it is unclear whether broad-specificity terminase recognition is unique to STAND-based sensors or shared by other defense systems. One enigmatic candidate is the defense protein Upx^17^, which contains a large, unannotated sensor domain and a PD-(D/E)xK nuclease domain but lacks a STAND NTPase. Although evolutionarily distinct from Avs proteins, the size of the Upx sensor suggests it may similarly recognize phage proteins. However, the specific triggers and the mechanism of Upx activation during infection remain unknown.

Here we combine cryo-electron microscopy (cryo-EM) with biochemical and functional analyses to define the basis of terminase recognition by Avs2. Furthermore, we show that Upx is also a broad-specificity terminase sensor, and structural modeling indicates that it binds an unfolded state of the terminase ATPase domain, an unexpected mode of phage protein recognition. Together, these results support terminase recognition as a conserved defense strategy and demonstrate how structurally diverse sensors converge on a shared viral target through distinct molecular mechanisms.

### Specificity of terminase sensing by Avs2

We previously showed that EcAvs2 confers defense against diverse types of tailed phages, including the T7-like phage PhiV-1 (ref.^17^) and the T1-like *Drexlerviridae* phage ZL19 (ref.^3^). We screened for EcAvs2 activators across the entire proteomes of these two phages using a co-expression toxicity assay, in which the expression of EcAvs2 with its cognate trigger in the same bacterial cell leads to EcAvs2-mediated cell death^8,17^ (Fig. 1a and Table S1). SeAvs3 was included in parallel as a control, as it confers defense against both PhiV-1 and ZL19 and is activated only by the terminase in each genome^3^. Our library included the complete genomes of PhiV-1 (49 genes) and ZL19 (77 genes), excluding four genes from each that were constitutively toxic to *E. coli* and not predicted to bind Avs2 or Avs3 by AlphaFold 3 (ref.^18^) (Table S2). For both phages, the large terminase was the only gene that induced EcAvs2- or SeAvs3-dependent toxicity (Fig. 1b). These results suggest that for these phages, the terminase is the PAMP recognized by EcAvs2 and SeAvs3, consistent with our previous findings^3^.

**Figure 1.**
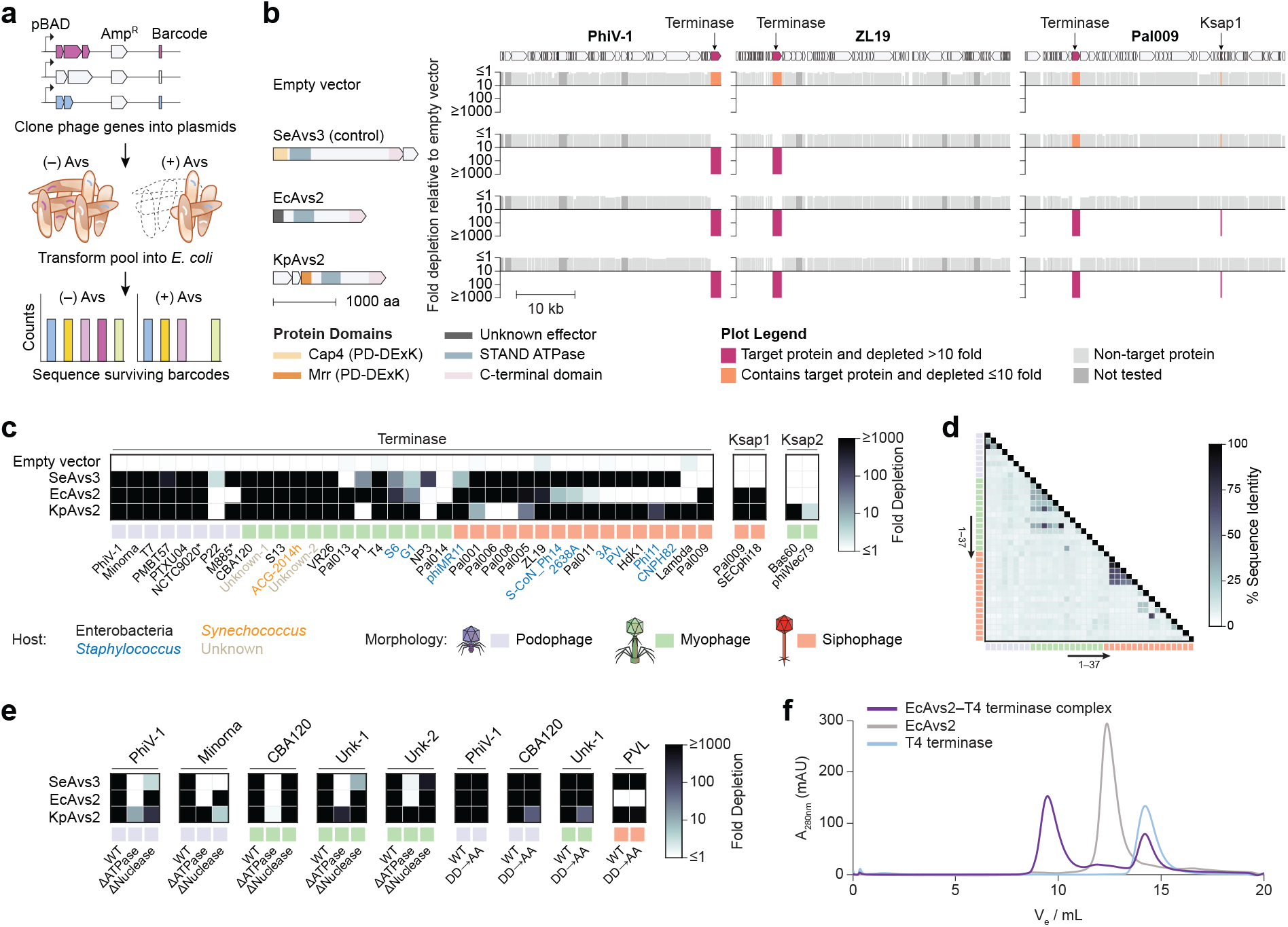
Avs2 is activated by diverse phage terminases. **a**, Schematic of the co-expression toxicity assay used to identify phage-encoded defense activators. **b**, Co-expression toxicity screen across the genomes of T7-like (PhiV-1), T1-like (ZL19), and *Dhillonvirus* (Pal009) phages. **c**, Heatmap of plasmid depletion for 37 terminases and Ksap1/2, organized by phage morphology. Asterisks indicate prophages. **d**, Percent amino acid identity matrix for the terminases evaluated in (c). **e**, Effect of terminase domain deletions and terminase active site mutations on co-expression toxicity. DD→AA refers to double alanine substitutions at the ATPase Walker B and nuclease catalytic aspartates (D161A/D365A, D357A/D499A, D233A/D384A, and D200A/D358A for phages PhiV-1, CBA120, Unk-1, and PVL, respectively). **f**, Size exclusion chromatography analysis of EcAvs2–T4 terminase complex formation.

An Avs2 homolog from *Klebsiella pneumoniae* (KpAvs2) can also be activated by two distinct proteins of unknown function, Ksap1 and Ksap2 (ref.^4^). These proteins are restricted to the *Dhillonvirus* genus and *Stephanstirmvirinae* subfamily, respectively, where they serve as the triggers of KpAvs2 during infection^4^. We identified a *Dhillonvirus* phage, Pal009, against which both EcAvs2 and KpAvs2 conferred robust defense (Fig. S1a). Mutation of the ATPase Walker A motif (K348A) of EcAvs2 ablated defense against Pal009 (Fig. S1a) as well as against T7 and PhiV-1 (Fig. S1b). To further investigate non-terminase activators, we expanded our library to include the complete Pal009 genome (58 genes), with the exception of one constitutively toxic gene that was not predicted to bind Avs2 or Avs3 by AlphaFold 3 (Table S2). The expanded library included Pal009 Ksap1, which shares 81% amino acid identity with the SECphi18 Ksap1 reported previously^4^. For both PhiV-1 and ZL19, KpAvs2 was toxic only in the presence of the terminase. However, for Pal009, both Avs2 homologs were toxic when co-expressed with either terminase or Ksap1, consistent with previous findings for KpAvs2 (ref.^4^) (Fig. 1b). By contrast, SeAvs3 was not activated by either Pal009 terminase or Ksap1, consistent with its lack of defense against Pal009 in plaque assays (Fig. S1a). While additional investigation is required to determine whether the Pal009 terminase or Ksap1 (or both) is the primary trigger of EcAvs2 during phage infection, these findings support co-expression toxicity as a robust strategy for identifying phage-encoded triggers of these defense systems.

To assess the generalizability of terminase detection, we constructed an expanded library of 37 diverse terminases, representing all three phage structural classes, along with two Ksap1 homologs from phages Pal009 and SECphi18 and two Ksap2 homologs from phages Bas60 and phiWec79 (Table S3). Consistent with previous observations, SeAvs3, EcAvs2, and KpAvs2 were toxic when co-expressed with a wide range of terminases (Fig. 1c). These terminases were highly divergent, with most sharing less than 10% pairwise sequence identity (Fig. 1d), consistent with the role of Avs proteins as pattern recognition receptors. Ksap1 activated both EcAvs2 and KpAvs2, whereas Ksap2 only activated KpAvs2 (Fig. 1c). Neither Ksap1 nor Ksap2 activated SeAvs3 (Fig. 1c).

Because the terminase consists of two distinct domains, an N-terminal ATPase and a C-terminal nuclease, we next investigated which of these domains is recognized by Avs2. We generated domain deletion variants of five terminases (PhiV-1, Minorna, CBA120, Unk-1, Unk-2) and tested them in coexpression toxicity assays. EcAvs2 was strongly activated by the ATPase domain of all five terminases (>1000-fold barcode depletion), whereas the nuclease domain exhibited little to no toxicity (Fig. 1e), in agreement with previous observations^3^. In contrast, KpAvs2 could be activated by both the ATPase and nuclease domains of Unk-1 and Unk-2, and by the nuclease domain of Minorna (Fig. 1e), suggesting that, at least in some cases, the nuclease domain of the terminase contributes to Avs2 recognition.

Finally, to investigate Avs2 activation biochemically, we analyzed recombinantly purified EcAvs2 and a representative terminase from phage T4 with size exclusion chromatography (SEC) in the presence of ATP and Mg^2+^. A high molecular weight peak appeared only when both proteins were present, indicating that EcAvs2 oligomerization is induced by terminase binding (Fig. 1f and Fig. S2).

### Cryo-EM structure of the EcAvs2−terminase complex

To investigate the mechanism of terminase recognition by Avs2, we assembled an EcAvs2−PhiV-1 terminase complex *in vitro* in the presence of ATP and Mg^2+^ and purified a stable complex by SEC (Fig. 2a). We determined the structure of this complex using single-particle cryo-EM. A reconstruction at 2.3 Å resolution revealed a C4-symmetric tetramer, consisting of four copies of the EcAvs2 protein and four copies of the PhiV-1 terminase (Fig. 2b and Fig. S3−4). This complex adopts a flat conformation, in contrast to the elongated tetrameric structure of terminase-bound SeAvs3 (ref.^3^) (Fig. S5a−b).

**Figure 2.**
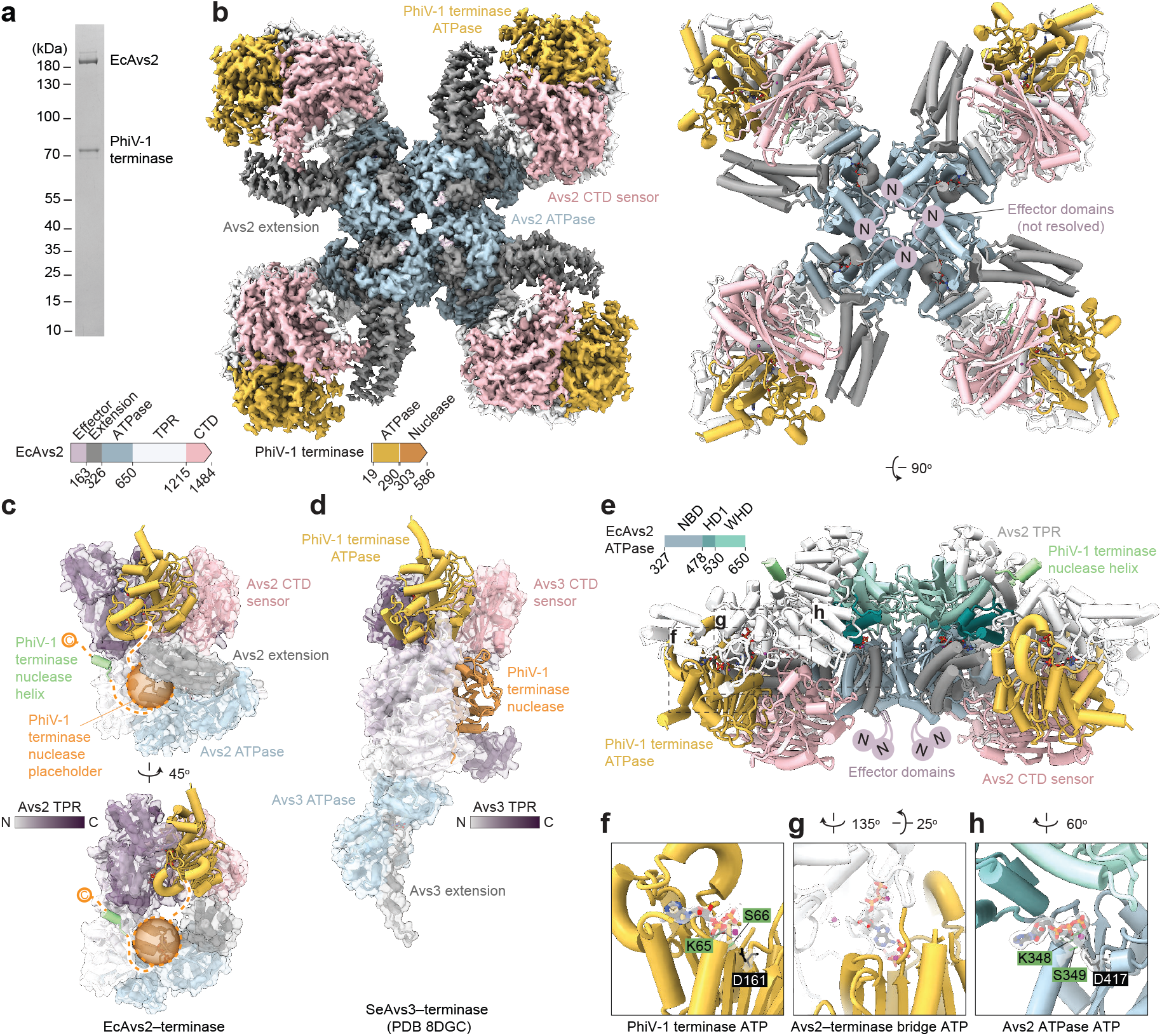
Cryo-EM structure of the EcAvs2–PhiV-1 terminase complex at 2.3 Å resolution. **a**, SDS-PAGE analysis of the purified EcAvs2– PhiV-1 terminase tetramer complex used for cryo-EM imaging. **b**, Cryo-EM density (left) and atomic model (right) of the tetramer, colored by domain. The EcAvs2 N-terminal effector domain is not visible in the cryo-EM density; its predicted location is indicated. **c, d**, Comparison between the protomers of the EcAvs2–terminase complex (**c**) and the SeAvs3–terminase complex (PDB 8DGC) (**d**). The EcAvs2 TPR domain is colored from N-to C-terminus as indicated. Green denotes the 11 resolved residues of the terminase nuclease domain; the putative position of the remainder of the nuclease domain is indicated by an orange sphere. **e**, Rotated view of the EcAvs2–terminase tetramer. The ATPase is colored by its subdomains: nucleotide-binding domain (NBD), helical domain 1 (HD1), and winged-helix domain (WHD). **f–h**, Insets show the ATP molecules and coordinating residues in the terminase ATPase (**f**), Avs2–terminase interface (**g**), and Avs2 ATPase (**h**). Green and black numbers indicate conserved residues within the Walker A and Walker B motifs, respectively.

The tetramerization interface of EcAvs2 is formed by its ATPase domain, each of which has an ATP molecule bound, consistent with the activated forms of other characterized STAND NTPases. Similar to SeAvs3, the portal-sensing EcAvs4 (ref.^3^), and the major capsid protein-sensing SeAvs7 (ref.^8^), tetramerization brings the N-termini of the EcAvs2 ATPase domains close together (Fig. 2b and Fig. S5a−b). In SeAvs3, EcAvs4, and SeAvs7, this proximity facilitates the tetramerization and activation of their N-terminal nuclease domains. In contrast, EcAvs2 has a distinct, uncharacterized effector domain with no clear enzymatic function or mechanism of toxicity. This effector was not visible in our cryo-EM reconstruction (Fig. 2b). Homologs of the effector are fused to diverse STAND NTPases and a novel PIN-domain containing system (Fig. S6a−b and Table S4), and AlphaFold 3 modeling suggests it might form a tetramer upon EcAvs2 oligomerization (Fig. S6c−e).

The remainder of EcAvs2 following the ATPase domain is distal to the symmetry axis and consists of an elaborate α-helical TPR domain, followed by a C-terminal β-strand-rich domain that we term the ‘C-terminal sensor.’ Together with a helical extension between the effector and ATPase domains, these form a large binding interface for the ATPase domain of the PhiV-1 terminase protein (Fig. 2b and Fig. S5a), burying 2833.5 Å^2^ of surface area. This interface is mediated by shape and charge complementarity (Fig. S7). The EcAvs2 TPR domain differs significantly from that of SeAvs3 (Fig. 2c−d and Fig. S5a−d), but both proteins share the C-terminal sensor, which binds to the same surface of the PhiV-1 termisnase ATPase (Fig. 2c−d). AlphaFold 3 models of EcAvs2 in complex with eight different phage terminases that exhibited co-expression toxicity (ipTM >0.7) are consistent with the EcAvs2−PhiV-1 terminase cryo-EM structure (Fig. S8), suggesting that this mode of terminase binding is conserved.

### Molecular basis of terminase recognition by EcAvs2

Our toxicity assays indicated that the nuclease domain of the PhiV-1 terminase is not required for EcAvs2 activation (Fig. 1e), and consistent with this, we did not observe cryo-EM density for most of the PhiV-1 terminase nuclease (Fig. 2c). However, we resolved a single 11-amino-acid helix from the nuclease domain lodged within a hydrophobic pocket on the EcAvs2 TPR domain (Fig. S5a,c−d and S8b). This interaction requires a large structural rearrangement of the natively folded terminase, as these residues are normally buried deep within the nuclease core (Fig. S5d), and superimposing the folded nuclease domain onto the EcAvs2-bound helix results in major steric clashes with the EcAvs2 TPR domain (Fig. S5e). These observations suggest that the terminase nuclease domain unfolds at least partially in order to bind to EcAvs2. This contrasts with SeAvs3, which extensively binds the terminase nuclease in its native conformation (Fig. 2d and Fig. S5b,d).

The EcAvs2−PhiV-1 terminase complex contains three ATP ligands (Fig. 2e−h), two of which are coordinated by canonical Walker motifs in the EcAvs2 and terminase ATPase domains, respectively. Notably, a third “bridging ATP” is sequestered by EcAvs2 to seal the inter-protein interface (Fig. 2g and Fig. 3a−b). This bridging ligand forms hydrogen bonds with both the γ-phosphate of terminase ATP molecule and the Mg^2+^ hydration shell of the terminase active site (Fig. 3a). EcAvs2 coordinates the bridging ATP through multiple contacts. Thr1128 and Arg1132 form H-bonds with the γ-phosphate of the bridging ATP, while Glu1364 forms H-bonds with its ribose (Fig. 3a−b). Asp981 forms a water-mediated hydrogen bond with the Mg^2+^ ion coordinated by the bridging ATP, as well as two direct hydrogen bonds with Arg1130, which coordinates the γ-phosphate of the terminase ATP (Fig. 3a).

**Figure 3.**
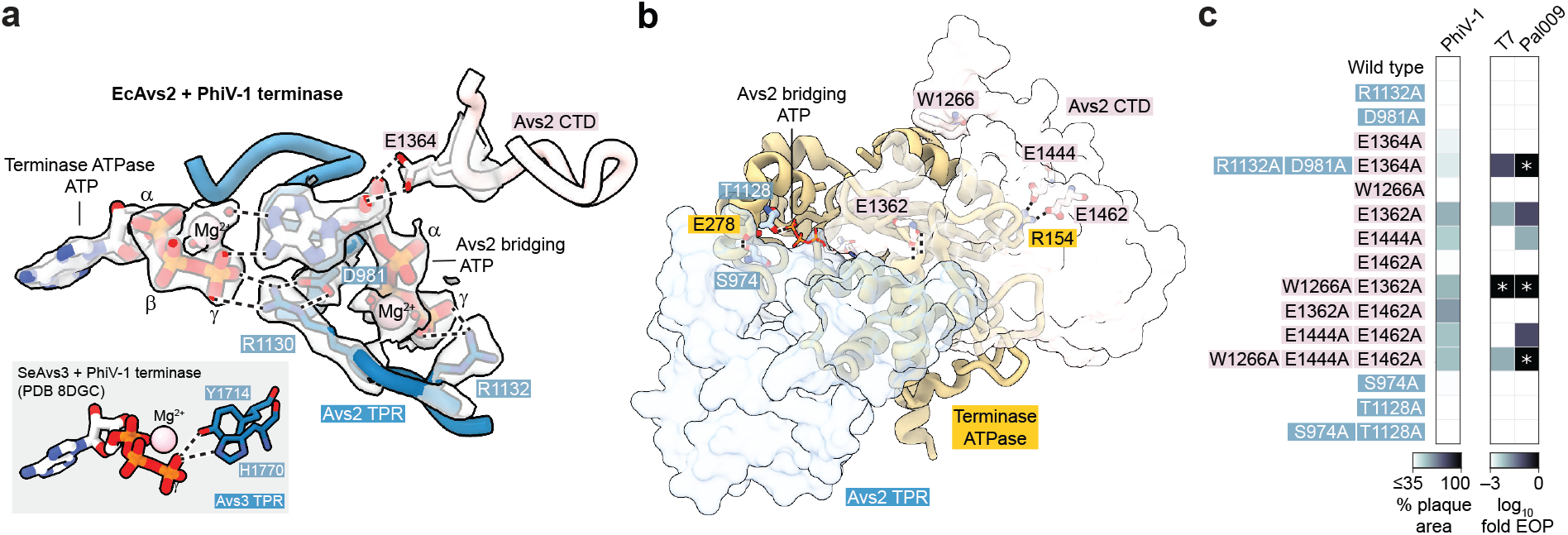
Combinatorial mutations in the EcAvs2 binding interface reduce defense activity. **a**, Interaction between the PhiV-1 terminase-bound ATP and a second ATP molecule bound by EcAvs2. Cryo-EM density is overlaid as a transparent grey surface. **b**, Additional interface contacts between EcAvs2 and the PhiV-1 terminase ATPase domain. EcAvs2 is shown as a transparent surface, with key interacting residues highlighted as sticks. **c**, Impact of single and combinatorial EcAvs2 mutations on phage plaque formation. Blue labels indicate residues interacting with the bridging ATP, and pink labels indicate residues interacting directly with the terminase. For PhiV-1, the heatmap represents percent plaque area relative to the empty vector control. For T7 and Pal009, the heatmaps show the log-fold reduction in efficiency of plating (EOP). Asterisks denote a reduction in plaque size compared to the empty vector control.

To test the importance of EcAvs2−terminase interactions, we introduced alanine substitutions in EcAvs2 targeting either the bridging ATP or the terminase interface and assayed phage defense against PhiV-1, T7, and Pal009. While individual alanine mutations of the bridging ATP-coordinating residues Arg1132, Asp981, Glu1364, or Thr1128 did not impair phage defense (Fig. 3c and Fig. S9a), the R1132A/D981A/E1364A triple mutation substantially disrupted defense against T7 and Pal009 (Fig. 3c and Fig. S9a). Many of the tested mutations disrupting residue−residue interactions also resulted in partial reduction of defense activity (Fig. 3b,c and Fig. S9a), including E1362A, which disrupts two hydrogen bonds with the backbone of the N-terminal base of a terminase alpha helix, and the E1444A/E1462A double mutant, which disrupts salt bridges with Arg154 of the terminase (Fig. 3c). The addition of W1266A, which stacks against a hydrophobic terminase surface (Trp179), to these two mutants enhanced their phenotypes. Western blotting confirmed that all EcAvs2 mutants retained wild-type level expression levels (Fig. S9b). Together, these results suggest that multiple redundant interactions contribute to terminase recognition by EcAvs2, consistent with its ability to detect diverse terminase proteins.

Although Ksap1 and Ksap2 are predicted to bind the same pocket of Avs2 as the PhiV-1 terminase, they interact primarily with the CTD sensor rather than the more extensive binding interface utilized by the terminase (Fig. 2c and Fig. S10). Consistent with this observation, the predicted buried surface area is substantially larger for Avs2−terminase than for Avs2−Ksap1/2 (e.g., 2833.5 Å^2^ for EcAvs2−terminase vs. 1573.6 Å^2^ for EcAvs2−Ksap1). Nevertheless, Ksap1 and Ksap2 adopt secondary structures similar to the terminase, featuring a central 4−6-stranded β-sheet and flanking helices that face the CTD sensor (Fig. S11).

To identify additional non-terminase activators, we performed an *in silico* AlphaFold 3 screen of EcAvs2 co-folded with individual proteins from phage T4 and a T4-like phage (PhiC120) (Fig. S12a−b). This screen identified vs.8, an uncharacterized ATPase-like protein, which was toxic when co-expressed with EcAvs2 in *E. coli* (Fig. S12c). vs.8 shares sequence homology with the ATPase domain of the T4 terminase (Fig. S12d−e), including conserved Walker A and Walker B motifs (Fig. S12e), and exhibits high structural similarity (Fig. S12f−h). An AlphaFold 3 model (EcAvs2−vs.8 ipTM 0.84) placed vs.8 within the EcAvs2 binding pocket at the same position as the PhiV-1 terminase ATPase domain, retaining the bridging ATP (EcAvs2−ATP ipTM 0.85) (Fig. S12i). Further investigation is required to determine the role of vs.8 in activating EcAvs2 during T4 infection.

### The defense protein Upx is a terminase sensor distinct from Avs1−3

We next investigated whether Upx, which has a large putative sensor domain, might also sense phage proteins. Upx is broadly distributed across bacterial and archaeal phyla and, in some cases, genomically associated with a split-TPR variant of the terminase-sensing Avs1 family (Fig. 4a, Fig. S13, and Table S5). Divergent Upx homologs from *Salmonella enterica* (SeUpx)^17^ and *Hydrobacter penzbergensis* (HpUpx) (sharing 16% amino acid identity) conferred robust defense against PhiV-1, consistent with previous findings for SeUpx^17,19,20^. HpUpx also conferred partial defense against the myophage T4 (Fig. 4b). Using the same genetic screen, we found that co-expression of both SeUpx and HpUpx with PhiV-1 genes led to the specific depletion of the PhiV-1 terminase, suggesting Upx also recognizes terminase to activate abortive infection or cellular toxicity. Both phage defense and terminase-induced toxicity were abolished in the SeUpx nuclease-dead mutant E1176A, within the ExK motif of its divergent C-terminal PD-(D/E)xK nuclease domain (Fig. 4b−c).

**Figure 4.**
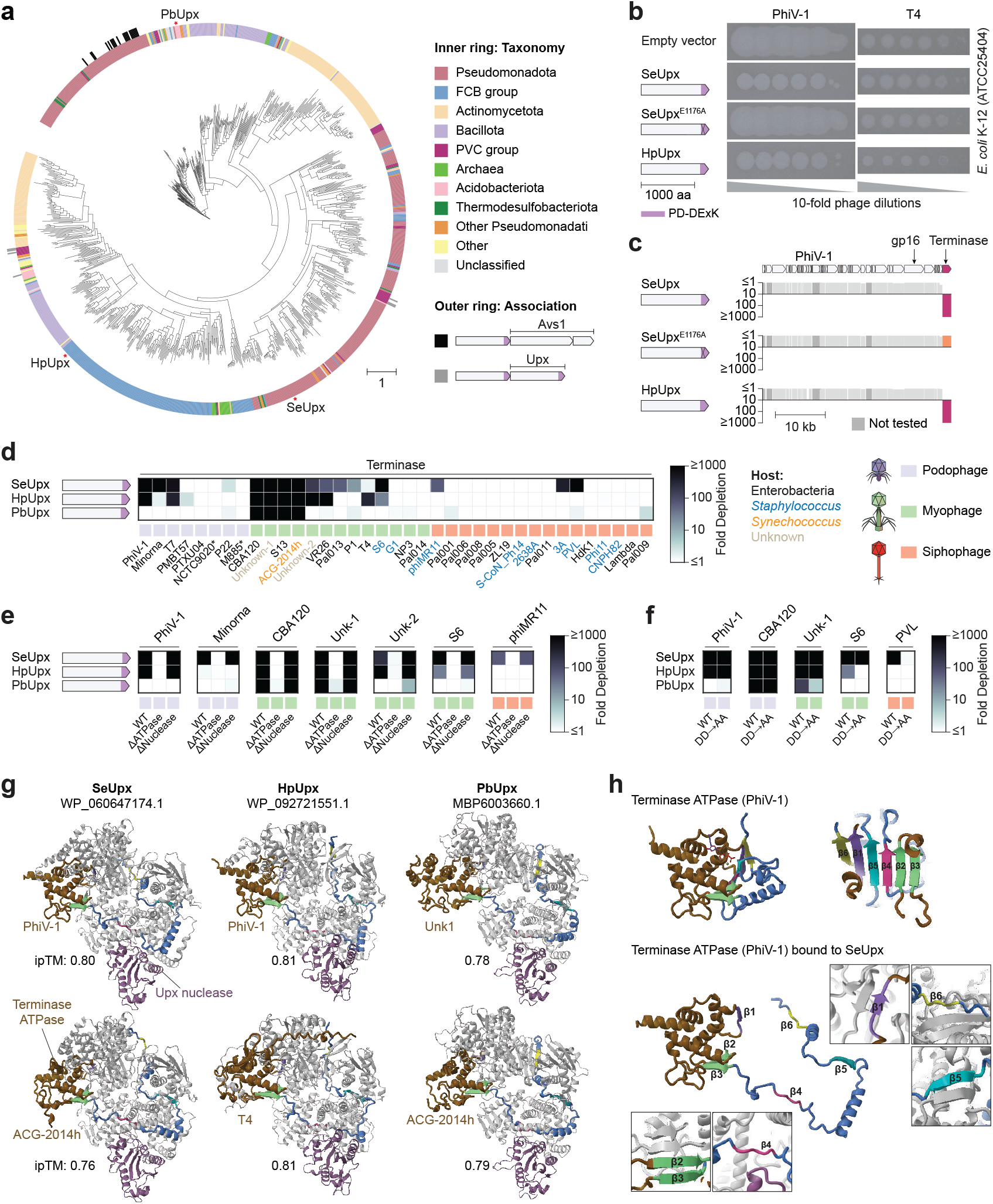
The bacterial defense protein Upx recognizes phage terminases. **a**, Maximum likelihood phylogenetic tree of 895 Upx representatives (clustered at 85% identity, 80% coverage). The inner ring indicates host phylum, and the outer ring indicates association with Avs1 or a second Upx. Red stars denote experimentally tested homologs from *Salmonella enterica* (SeUpx), *Hydrobacter penzbergensis* (HpUpx) and a *Pyrinomonadaceae* bacterium (PbUpx). **b**, Defense activity of SeUpx, HpUpx, and PbUpx against phages PhiV-1 and T4 in *E. coli* K-12 (ATCC 25404). Spots correspond to 10-fold serial dilutions of phage from left to right. **c**, Co-expression toxicity screen of SeUpx, the nuclease-dead SeUpx^E1176A^ mutant, and HpUpx across the PhiV-1 genome. **d**, Heatmap of plasmid depletion for 37 terminases. Asterisks indicate prophages. **e, f**, Effect of terminase domain deletions (**e**) and terminase active site mutations (**f**) on Upx-mediated toxicity. DD→AA refers to double alanine substitutions at the ATPase Walker B and nuclease catalytic aspartates (D225A/D415A for phage S6; see also Fig. 1 legend). **g**, AlphaFold 3 models of SeUpx, HpUpx, and PbUpx in complex with terminase ATPase domains (ligand-free). ipTM, interface predicted template modeling score. **h**, Structural comparison of the PhiV-1 terminase ATPase alone (top) versus the SeUpx-bound state (bottom), suggesting SeUpx recognizes an unfolded ATPase conformation.

To interrogate recognition specificity, we assessed the coexpression toxicity of SeUpx, HpUpx, and an additional homolog from a *Pyrinomonadaceae* bacterium (PbUpx). All three were activated by a diverse set of terminases, spanning all three phage structural classes (Fig. 4d). The recognized terminases have limited homology, with some pairs sharing <10% amino acid identity. Terminase truncations showed that the ATPase domain alone is sufficient to activate all Upx homologs (Fig. 4e). In addition, double alanine substitutions of the conserved aspartates in the ATPase Walker B and nuclease catalytic sites of the terminase preserved Upx-mediated toxicity in most cases (Fig. 4f), indicating that Upx does not depend on terminase catalytic activity.

High-confidence AlphaFold 3 models of Upx−terminase complexes (ipTM ~0.8) suggest an unexpected recognition mode in which Upx binds a locally unfolded state of the terminase ATPase domain (Fig. 4g). In these models, several β-strands that normally pack together to form the ATPase core β-sheet (β1, β4, β5, and β6) are disengaged and instead threaded through the Upx N-terminal sensor domain. Several unfolded β-strands are predicted to form backbone hydrogen bonds with Upx β-strands through β-augmentation (Fig. 4h and Fig. S14), a mode of recognition that does not depend on amino acid sequence. Notably, this unfolded ATPase conformation and its placement through Upx are predicted consistently across six diverse terminases and across three Upx homologs (Fig. 4g−h, Fig. S14c, and Fig. S15), suggesting a conserved binding geometry despite limited terminase sequence similarity. Finally, a model of PbUpx co-folded with full ACG-2014h terminase suggests the terminase nuclease domain does not substantially interact with Upx (Fig. S14e and Fig. S15).

## Discussion

Here, we describe the molecular basis of phage large terminase subunit recognition by the bacterial defense proteins Avs2 and Upx. These terminase sensors reveal unexpected mechanisms of phage protein recognition, including utilizing an ATP ligand to facilitate an interaction interface and predicted binding to unfolded target states.

The EcAvs2 complex is structurally distinct from other characterized STAND NTPases. EcAvs2 forms a flat tetramer in which each protomer contains a large helical extension between the N-terminal effector and the ATPase domain that is proximal to the terminase, as well as an unexpected bridging ATP that directly interacts with both protein residues and the canonical terminase ATP. Multiple electrostatic and hydrophobic interactions at the binding interface provide functional redundancy, possibly enabling EcAvs2 to detect highly diverse terminases with minimal sequence similarity. These interactions might also contribute additively to a conformational rearrangement or rigidification of the TPR domain upon terminase binding, transmitting an allosteric signal to the ATPase that promotes oligomerization^3,8,21,22^.

While most Avs2 homologs utilize Mrr-like dsDNA nucleases as effectors^3^, the function of the less common EcAvs2 effector remains unknown. This domain appears to be highly modular, occurring at the N-terminus of at least eight other defense systems (Fig. S6a−b, Table S4). Its first 15 residues are homologous to amyloid signaling proteins involved in cell death^23^, and its sequence contains conserved non-polar residues but lacks conserved charged or polar residues (see Supplementary Data), suggesting a non-enzymatic function. Consistent with this, we observed no cryo-EM density for the effector in the terminase-bound tetramer, indicating it is disordered or flexible in the absence of membranes or interacting partners. Further work such as affinity purification and mass spectrometry of EcAvs2 during phage infection, or the isolation of phage escape mutants and host mutants resistant to EcAvs2 toxicity, is required to determine its activity and the requirement for oligomerization.

Although non-terminase activators such as Ksap1, Ksap2, and possibly vs.8 are intriguing, several lines of evidence suggest that the terminase is the primary evolutionary target of Avs2. The Avs2−terminase complex is defined by strong shape and charge complementarity between the Avs2 sensor and the terminase. While Ksap1, Ksap2, and vs.8 share structural similarity with the terminase ATPase and are predicted to bind the Avs2 pocket in the same orientation, they do not engage the full binding interface, suggesting that Avs2 is adapted for the terminase. Consistent with this hypothesis, Avs2 appears to have a secondary binding interface that can in some cases interact with the terminase nuclease domain, an interaction that is absent with the three non-terminase activators. Moreover, Ksap1, Ksap2, and vs.8 are restricted to specific phage taxa, whereas the terminase is universal across tailed phages, consistent with the broad anti-phage specificity of Avs2 and its presence across more than 20 prokaryotic phyla^3^. Together, these observations support a model in which the terminase is the canonical trigger of Avs2, with non-terminase activators representing lineage-specific exceptions.

The function of these non-terminase activators remains an interesting open question, as it is unlikely that they exist primarily to trigger Avs2, given that this would be counterproductive for the phage life cycle. One possibility is that they function as anti-defense proteins that Avs2 evolved to recognize as additional triggers. Examples of this phenomenon include the PARIS and Ronin defense systems, which are activated by the anti-restriction proteins Ocr and ArdA, respectively^24,25^. Further investigation is required to determine the role of these proteins in the phage life cycle.

While recent work proposed the PhiV-1 ejection protein gp16 as a trigger for SeUpx^19^, our results indicate that Upx senses the large terminase (gp19 in PhiV-1). In a systematic toxicity screen of the PhiV-1 genome, gp19 emerged as the sole activator of both SeUpx and HpUpx, whereas gp16 failed to induce toxicity. Furthermore, divergent terminase proteins strongly activated three distinct Upx homologs in toxicity assays, and HpUpx also conferred partial defense against phage T4, a myophage that lacks a gp16 homolog. The potent toxicity induced by these terminases suggests that Upx, like Avs2, acts through programmed cell death to halt infection. Finally, AlphaFold 3 models of diverse Upx−terminase complexes exhibited a consistent orientation of the unfolded terminase ATPase domain across all homologs, suggesting that this mode of terminase recognition is conserved across the Upx family. As with Avs2, however, our findings do not exclude the possibility that Upx may recognize additional non-terminase activators (e.g., ATPases) in other phage lineages.

The predicted mechanism of sensing by Upx, via recognition of a locally unfolded state of a globular protein, represents a departure from typical protein-protein interactions in immunity. This mechanism, supported by β-augmentation, might allow Upx to bypass sequence-level divergence and detect conserved structural features embedded within a core fold. It remains an open question whether Upx actively unfolds a native terminase ATPase domain, or might only bind nascent terminases co-translationally. Additionally, while Upx is a monomer in its apo form^19,20^, further investigation is required to determine whether terminase binding induces Upx oligomerization, similar to Avs2.

Altogether, our structural characterization of EcAvs2 and identification of Upx as a terminase sensor support the terminase as a key trigger of bacterial defense. The recognition of targets through ligand-mediated interfaces and binding of partially unfolded states represent mechanistic innovations that expand the repertoire of strategies bacteria use to detect phage infection.

## Materials and Methods

### Cloning

Genes were chemically synthesized (Twist Bioscience or Integrated DNA Technologies) or amplified by PCR from phage genomic DNA. Plasmids were constructed with Gibson assembly, purified using Qiagen miniprep buffers and 96-well DNA & RNA binding plates (Epoch Life Science), and fully sequence verified by Tn5 tagmentation and deep sequencing, as previously described^17,26^. Phage genes were cloned into barcoded pBAD expression vectors with ampicillin resistance. Defense genes were cloned with their native promoters and coding sequences into a pACYC184-like vector backbone with chloramphenicol resistance.

### Competent cell production

*E. coli* strains were cultured in ZymoBroth with 25 µg/mL chloramphenicol and made competent using a Mix & Go Transformation Kit (Zymo).

### Co-expression toxicity screens

Barcoded pBAD vectors containing phage genes were pooled to generate a phage gene library with a balanced representation of genes. Control plasmids expressing mNeonGreen were then added to the library as normalization controls. The resulting gene library was transformed into *E. coli* NovaBlue(DE3) cells expressing defense systems of interest. After 1 hr outgrowth at 37 °C in SOC, cells were plated on LB agar containing 100 µg/mL carbenicillin, 25 µg/mL chloramphenicol, and 0.05% arabinose. Cells were further incubated for 12 h at 37 °C, and plasmids from surviving colonies were subsequently isolated by miniprep. Barcodes from these plasmids were amplified and sequenced on an Illumina NextSeq 550. Fold depletion values were calculated as previously described^8^.

### Cryo-EM sample preparation

EcAvs2 and PhiV-1 terminase gene sequences were codon optimized for expression in *E. coli* K-12. EcAvs2 was cloned into a pCDF plasmid with a C-terminal TwinStrep tag. PhiV-1 terminase was cloned into a pBR322 plasmid with a C-terminal 6xHis tag. EcAvs2 and PhiV-1 were expressed in *E. coli* BL21(DE3) cells. EcAvs2 cultures were grown at 37 °C in Terrific Broth (TB) containing 50 µg/mL spectinomycin, induced with 0.25 mM isopropyl β-D-1-thiogalactopyranoside (IPTG) at an OD_600_ of approximately 0.5, and subsequently grown at 18 °C overnight.

PhiV-1 terminase cultures were grown in TB containing autoinduction buffer^27^ (25 mM Na_2_HPO_4_, 25 mM KH_2_PO_4_, 50 mM NH_4_Cl, 5 mM Na_2_SO_4_, 0.5% w/v glycerol, 0.2% w/v α-Lactose monohydrate, 0.05% w/v glucose, and 2 mM MgSO_4_) and 100 µg/mL ampicillin at 37 °C for ~8 h followed by 18 °C overnight. Cells were harvested by centrifugation and resuspended in lysis buffer containing 50 mM Tris-HCl (pH 7.4), 500 mM NaCl, 5% v/v glycerol and 5 mM 2-mercaptoethanol. For PhiV-1 terminase, the lysis buffer was supplemented with 20 mM imidazole. After lysis via an LM20 Microfluidizer (Microfluidics) and clarification by centrifugation, the supernatants for EcAvs2 and PhiV-1 terminase were incubated with Strep-Tactin Superflow Plus (Qiagen) or Ni-NTA Superflow resin (Qiagen), respectively. Resin-bound proteins were washed with their respective lysis buffer and eluted with lysis buffer containing 5 mM desthiobiotin (for EcAvs2) or 300 mM imidazole (for PhiV-1 terminase). Eluted proteins were concentrated using Amicon Ultra centrifugal filters (Sigma Aldrich) with molecular weight cutoffs of 50 kDa for EcAvs2 and 10 kDa for PhiV-1 terminase, yielding final concentrations of 17 mg/mL and 12 mg/mL, respectively.

The EcAvs2-terminase complex was assembled by incubating EcAvs2 (~10 µM) with PhiV-1 terminase (~70 µM) in a buffer containing 25 mM Tris-HCl (pH 7.4), 250 mM NaCl, 2.5% v/v glycerol, 2.5 mM 2-mercaptoethanol, 1 mM ATP, and 5 mM MgCl_2_. The reaction was carried out at 37 °C for 1 hour. Precipitates were pelleted by centrifugation, and the soluble complex in the supernatant was purified using a Superose 6 Increase 10/300 GL column with a buffer containing 20 mM Tris-HCl (pH 7.4), 200 mM NaCl, 0.1 mM ATP, and 2 mM MgCl_2_. Fractions containing the oligomeric EcAvs2-terminase complex were pooled and concentrated to 1.8 mg/mL using a 50 kDa Amicon Ultra centrifugal filter.

For cryo-EM grid preparation, a freshly glow-discharged (60 s at 25 mA) Cu300 R1.2/1.3 holey carbon grid (Quantifoil) was mounted in the chamber of a Vitrobot Mark IV (Thermo Fisher Scientific) maintained at 12 °C and 100% humidity. A 4 µL aliquot of EcAvs2−PhiV1 terminase complex was applied to the grid, immediately blotted using Ø55 grade 595 filter paper (Ted Pella), and plunged into liquid ethane.

### EcAvs2−T4 terminase complex analysis

The codon-optimized sequence for EcAvs2 and the native sequence for T4 termi-nase were individually cloned into a pSF1389 plasmid^28^ with an N-terminal 14xHis tag followed by a brNedd8 cleavage site. Each protein was expressed in *E. coli* NEBExpress (New England Biolabs). Cultures were grown at 37 °C in Super Broth (SB) supplemented with 50 µg/mL kanamycin, induced with 0.2 mM IPTG at an OD_600_ of approximately 0.8, and subsequently grown at 18 °C overnight. Cells were harvested by centrifugation and resuspended in lysis buffer containing 50 mM Tris-HCl (pH 7.5), 300 mM NaCl, 20 mM imidazole, 1 mM DTT, and 1 mM phenylmethylsulfonyl fluoride (PMSF). Following lysis by sonication and clarification by centrifugation, the supernatant was incubated with Ni-NTA Superflow resin (Qiagen). The resin was washed with lysis buffer, and the proteins were eluted with lysis buffer supplemented with 300 mM imidazole. The eluted protein was concentrated using an Amicon Ultra centrifugal filter (Sigma Aldrich) with a 10 kDa molecular weight cutoff for T4 terminase and a 100 kDa molecular weight cut off for EcAvs2. Proteins were further purified via size exclusion chromatography using a Superdex 200 Increase 10/300 GL column pre-equilibrated with 50 mM HEPES (pH 7.5), 250 mM NaCl, 2 mM Mg(OAc)_2_, 0.1 mM ATP, and 1 mM DTT. Pure fractions were analyzed by SDS-PAGE, pooled, concentrated to ~10 mg/mL, and stored at −80 °C.

Purified EcAvs2 and T4 terminase were mixed at a 1:2 molar ratio to assemble the EcAvs2−T4 terminase complex. The individual proteins and assembled complex were then analyzed by size exclusion chromatography. For each run, a 450 µL sample containing ~0.5−1.0 mg of total protein was loaded onto a Superdex 200 Increase 10/300 GL column (Cytiva) and eluted with a buffer containing 50 mM HEPES (pH 7.5), 250 mM NaCl, 2 mM Mg(OAc)_2_, 0.1 mM ATP, and 1 mM DTT.

### Terminase alignment generation

A structure-guided multiple sequence alignment (MSA) of the 37 terminases was generated using a hierarchical approach. Individual structural models were first predicted using AlphaFold 3 (ref.^18^). Sequences were then partitioned into six groups based on sequence similarity and aligned using PROMALS3D^29^. These sub-alignments were integrated into a global MSA using an additional PROMALS3D alignment of group representatives as a template. The global MSA was then trimmed to the core terminase domain to calculate percent amino acid identity values.

### Cryo-EM data collection

Cryo-EM data were collected using the Thermo Scientific Titan Krios G3i cryo-TEM at MIT.nano using a K3 direct detector (Gatan) operated in super-resolution mode with 2-fold binning, and an energy filter with slit width of 20 eV. Micrographs were collected automatically using EPU in AFIS mode, yielding 16,853 movies at 130,000× magnification with a real pixel size of 0.663 Å. The defocus range was set from −1.8 µm to −0.8 µm with an exposure time of 0.52 s and fractionated into 32 frames at a flux of 27.4 e^−^/pix/s, resulting in a total fluence per micrograph of 32.4 e^−^/Å^2^.

### Cryo-EM data processing

Cryo-EM data were processed using RELION^30^. Movies were corrected for motion using the RELION implementation of MotionCor2 (ref.^31^), with 5 × 5 patches and dose-weighting. CTF parameters were estimated using CTFFIND-4.1 (ref.^32^). Particle picking was performed using Topaz with the pre-trained general model^33^, yielding 2,840,455 particles that were extracted in a 400 Å box down-sampled to 4 Å/pix. A subset of 177,451 particles selected by 2D classification were used to generate an ab initio 3D volume with C1 symmetry, to which C4 symmetry was subsequently applied. The symmetrized map was used as a reference for 3D classification of all particles, with C1 symmetry applied during classification. A subset of 1,416,520 particles (50%) was selected and refined with C4 symmetry to the downsampled Nyquist limit of 8 Å resolution. To select a higher-quality subset, 3D classification was performed without alignment using a 260 Å diameter circular mask, with C4 symmetry applied and a regularization parameter (T) of 8, yielding 353,169 particles. These particles were re-extracted and refined with C4 symmetry and masking to 2.95 Å resolution. Particles were further improved by Bayesian polishing (with parameters determined by training s_vel: 0.942, s_div: 15060, s_acc: 1.815) with polished particles extracted with 0.9945 Å/pix and a 400-pixel box (rescaled from 600 pixels). After successive rounds of CTF refinement, correcting for anisotropic magnification, per-particle defocus, beam tilt, trefoil, and 4th-order aberrations, the particles were refined with C4 symmetry to 2.33 Å resolution.

Close to the symmetry axis, this reconstruction exhibited density features consistent with the estimated resolution, including well-defined carbonyl atoms and clear solvent density. However, resolution decreased further from the axis near the terminase protein, indicative of imperfect symmetry. The particles were therefore subjected to C4 symmetry expansion, followed by refinement with C1 symmetry with a soft mask around only the terminase and contacting regions, with local angular sampling starting at 1.8°. This produced a reconstruction with 2.23 Å resolution (within the refinement mask) with clearer features further from the symmetry axis. Resolution is reported using the gold-standard Fourier Shell Correlation with 0.143 cutoff.

### Model building and refinement

For model building, the focus-refined map was symmetrized and then combined with the overall map using the ‘vol max’ command in UCSF ChimeraX^34^ to produce a composite map. A model of EcAvs2 generated with AlphaFold2 was docked into the density and manually adjusted, first with ChimeraX and subsequently in Coot^35^. The structure of PhiV-1 terminase was docked from our previous cryo-EM structure of PhiV-1 terminase in complex with SeAvs3. All residues were inspected manually in Coot and adjusted or removed as necessary. Ordered water molecules were automatically placed into >9σ density peaks using Coot, followed by manual curation to ensure reasonable coordination distances. The model was symmetrized and then refined using ISOLDE^36^. A single asymmetric unit was manually fixed prior to re-symmetrization and refinement using phenix.real_space_refine^37^ with starting model restraints. Structural figures were generated using ChimeraX.

### Plaque assays

A total of 1 mL of *E. coli* strain C expressing a defense system, mutant, or empty vector were grown in Terrific Broth to an OD_600_ of 0.8−1.0 and added to 49 mL top agar (7 g/L agar, 15.5 g/L LB Lennox, 4.5 g/L NaCl) with 25 µg/mL chloramphenicol, poured into a 15 cm Petri dish, and allowed to solidify at room temperature. Ten-fold dilutions of phage in deionized water were spotted on top, with 3 µL per spot. After a 17-hour incubation at 37 °C, plates were imaged in the dark over a white backlight using a Nikon D7500 DSLR camera.

### Western blotting

C-terminal TwinStrep tags were cloned into the constructs for wild-type and mutant EcAvs2. Proteins were expressed in *E. coli* strain C. For each protein, a total of 250 mL of TB containing 25 µg/mL chloramphenicol was seeded with overnight culture and shaken at 37 °C for 14 h. Cells were harvested by centrifugation and resuspended in lysis buffer containing 50 mM Tris-HCl (pH 7.4), 250 mM NaCl, 5% v/v glycerol and 5 mM 2-mercaptoethanol. Cells were lysed using an LM20 Microfluidizer at 28,000 PSI, and lysates were clarified by centrifugation. The resulting supernatants were incubated with 300 µL of Strep-Tactin Superflow Plus resin (Qiagen) and rotated at 18 °C for 1 h. The suspension was poured directly into a 10 mL gravity flow column, using additional lysis buffer to collect any residual resin. Resin-bound proteins were washed with lysis buffer and eluted with lysis buffer containing 5 mM desthiobiotin.

For immunoblotting, eluted samples were resolved on a 4−12% precast polyacrylamide gel (Invitrogen) at 4 °C for 3 h and transferred onto nitrocellulose membranes using a TransBlot Turbo Transfer System (Bio-Rad). The membrane was blocked for 1 h in 5% non-fat milk with TBST, followed by overnight incubation at 4 °C with a 1:1000 dilution of Strep-Tag II monoclonal primary antibody (Invitrogen). After washing, membranes were incubated with a 1:5000 dilution of a horseradish peroxidase (HRP)-conjugated anti-mouse secondary antibody (Jackson ImmunoResearch) for 1 h at room temperature. Protein bands were detected using Pierce ECL Western Blotting Substrate (Thermo Fisher Scientific) and visualized on film using an SRX-101A film processor.

### Upx phylogeny

Upx homologs (Table S5) were identified through PSI-BLAST^38^ searches against the NCBI nonredundant (nr) protein sequence database (December 2024). Searches were seeded on two subsequences within a representative Upx homolog (WP_141688245.1), corresponding to residues 50−461 and 502−916, and iterated until convergence. For each iteration, hits were required to have an *E*-value below 0.001 and at least 50% coverage. Following removal of truncated proteins, this procedure yielded 2,084 hits.

To identify additional Upx homologs potentially missed during the initial searches, a multiple sequence alignment (MSA) of representatives from the 2,084 hits was first generated using MAFFT^39^. A position-specific scoring matrix (PSSM) was then constructed for a conserved central region (residues 668−1067 in WP_060647174.1) using RPSBLAST. This PSSM was used to perform a final PSI-BLAST search against nr, requiring a maximum *E*-value of 0.01 and at least 50% coverage, which identified 9 additional sequences.

The complete set of 2,093 sequences was clustered using MM-seqs2 (ref.^40^) at 85% sequence identity and 80% coverage, resulting in 895 clusters. A single representative sequence from each cluster was selected for phylogenetic analyses. An MSA of the representative sequences was generated using MAFFT v7.520 (ref.^39^) with global pairwise alignment (--maxiterate 1000 --globalpair) and trimmed with trimAl v1.2 (ref.^41^) using a gap threshold of 0.25 (-gt 0.25). A maximum likelihood phylogenetic tree was constructed from the trimmed alignment using IQ-TREE v1.6.12 (ref.^42^) with the LG+G4 model and 2000 ultrafast bootstrap replicates (-nstop 500 -safe -nt 4 -bb 2000 -m LG+G4). Taxonomic classifications were retrieved from the NCBI taxonomy database, and the tree was visualized using iTOL^43^.

### EcAvs2 effector domain phylogeny

EcAvs2 effector domain homologs were retrieved using a PSI-BLAST search against nr (March 2025), starting with a custom position-specific scoring matrix (PSSM) for the effector domain. Hits were required to have an *E*-value below 0.01. Sequences were clustered using MMseqs2 at 95% identity. Following removal of truncated proteins, 160 clusters (containing a total of 556 unique proteins) were retained for subsequent analyses. One representative from each cluster was selected, and an MSA of the representative sequences was generated using MAFFT v7.520 with global pairwise alignment (--maxiterate 1000 --globalpair). A maximum likelihood phylogenetic tree was constructed from the MSA using IQ-TREE v1.6.12 with the LG+G4 model and 2000 ultrafast bootstrap replicates (-nstop 500 -safe -nt 4 -bb 2000 -m LG+G4). Taxonomic classifications w ere retrieved from the NCBI taxonomy database, and the tree was visualized using iTOL.

### vs.8 phylogeny

Homologs of vs.8 were retrieved by performing five i terations o f P SI-BLAST a gainst n r (October 2025) with an initial *E*-value threshold of 0.005 and a final cutoff of 10^−30^. This search identified both vs.8 homologs and T4-like terminases, totaling 1,240 sequences after the removal of partial and truncated proteins. To reduce redundancy, sequences were clustered using MMseqs2 at 95% identity and 90% coverage (--min-seq-id 0.95 -c 0.9), resulting in 419 representatives. A structure-guided MSA was constructed by first aligning 10 representatives with PROMALS3D and subsequently adding the remaining 409 sequences using the --seed option in MAFFT. A maximum likelihood phylogenetic tree was constructed from the MSA using IQ-TREE v1.6.12 with the LG+G4 model and 2000 ultrafast bootstrap replicates (-nstop 500 -safe -nt 4 -bb 2000 -m LG+G4).

## ACKNOWLEDGEMENTS

We thank Ed Brignole and Chris Borsa for the smooth running of the MIT.nano cryo-EM facility, established in part with financial support from the Arnold and Mabel Beckman Foundation; and the Gao lab for insight and advice. C.C. was supported by an NIH Cellular and Molecular Training Grant (NIGMS, grant number 5T32GM007276). S.A.E. was supported by Stanford Bio-X Bowes fellowship and an NIH Genetics and Developmental Biology Training Grant (NIGMS, grant number 5T32GM141828). M.E.W. was supported by a postdoctoral fellowship from the Helen Hay Whitney Foundation. M.B. and T.P. were supported by Stanford Bio-X. F.Z. was supported by the Howard Hughes Medical Institute; Yang Tan Collective at MIT, the K. Lisa Yang and Hock E. Tan Molecular Therapeutics Center; Broad Institute Programmable Therapeutics Donors; and the BT Charitable Foundation. A.G. was supported by the G. Harold & Leila Y. Mathers Foundation, the Esther Ehrman Lazard Faculty Scholars Program, and Stanford Bio-X.

## AUTHOR CONTRIBUTIONS

S.A.E., C.C., and D.L. performed co-expression toxicity experiments. S.A.E., D.L., M.B., J.S., and A.G. purified proteins. M.B. performed size exclusion chromatography analysis. A.G. prepared the cryo-EM sample. M.E.W. collected cryo-EM data and determined structures. C.C., S.A.E., and A.G. generated AlphaFold models. C.C. and S.A.E. performed phage plaque assays. S.A.E. performed Western blot analysis. S.A.E. and A.G. performed phylogenetic analyses. All authors analyzed data. T.P., F.Z., and A.G. supervised research. S.A.E., C.C., M.E.W., and A.G. wrote the manuscript with input from all authors.

## COMPETING INTERESTS

F.Z. is a scientific advisor and cofounder of Beam Therapeutics, Pairwise Plants, Arbor Biotechnologies, Aera Therapeutics, and Moonwalk Biosciences. F.Z. is a scientific advisor for Octant. The remaining authors have no competing interests to declare.

## DATA AVAILABILITY

Cryo-EM maps have been deposited in the Electron Microscopy Data Bank with accession codes EMD-70096 and EMD-70097. The corresponding atomic model has been deposited in the Protein Data Bank under the accession code 9O4E

