## Supplementary Figures 1-15 for "Phage terminase recognition by the bacterial immune sensors Avs2 and Upx"

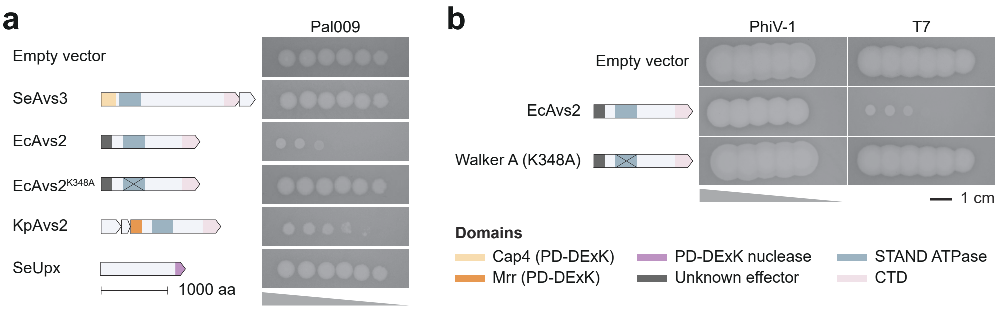


**Supplementary Fig. 1 | Anti-phage activity of SeAvs3, EcAvs2, KpAvs2, and SeUpx. a, b**, Plaque assays of indicated defense systems against *Dhillonvirus* phage Pal009 (**a**) and phages PhiV-1 and T7 (**b**). K348 is located in the Walker A motif of the EcAvs2 ATPase. Spots correspond to 10-fold serial dilutions of phage from left to right.

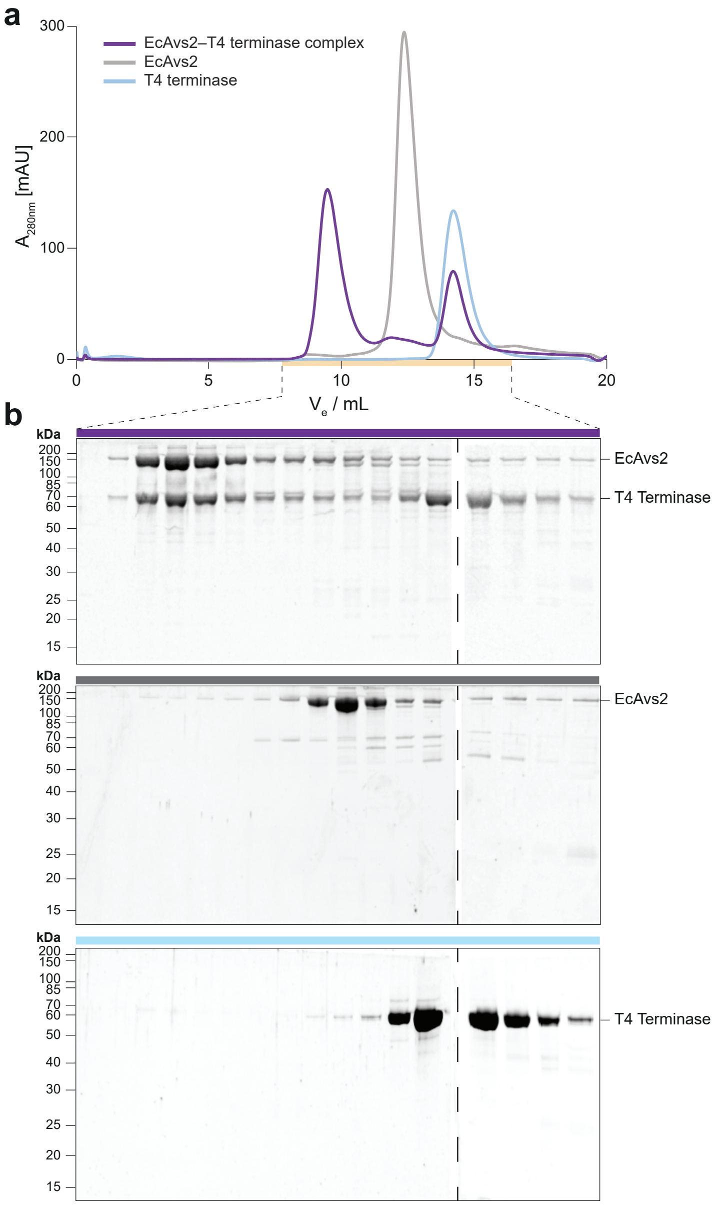


**Supplementary Fig. 2 | EcAvs2 oligomerizes in response to terminase binding. a**, Size exclusion chromatography profiles of EcAvs2, T4 terminase, and their mixture. **b**, SDS-PAGE analysis of the elution fractions indicated by the orange line in (a).

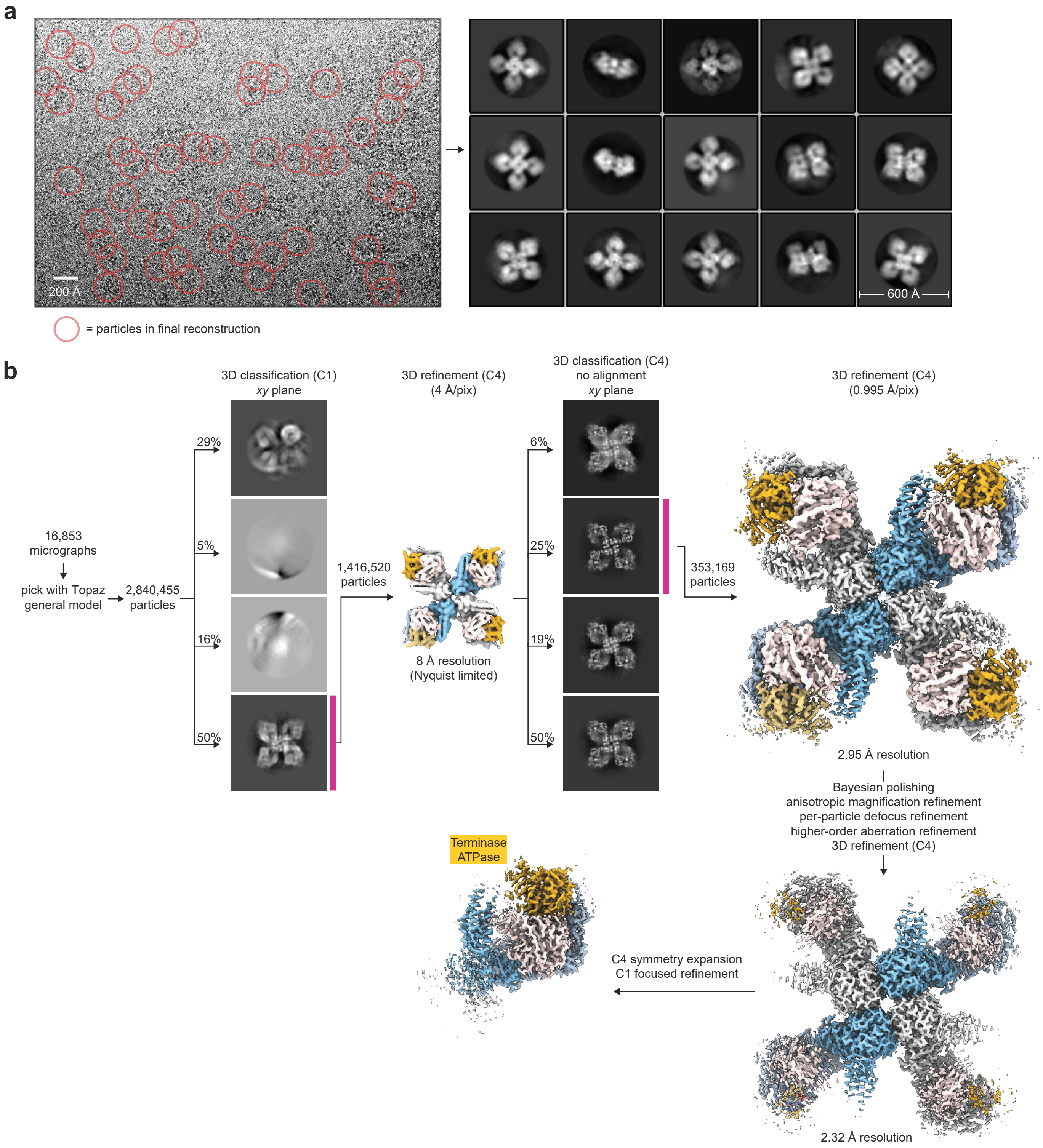


**Supplementary Fig. 3 | Cryo-EM image processing for the EcAvs2–PhiV-1 terminase complex (related to Fig. 2). a**, Left: Representative raw cryo-EM micrograph with particles in the final reconstruction circled. Right: Representative 2D class averages of the initially picked particles. 2D classification was not used to select particles in the final reconstruction. **b**, Flowchart of the RELION data processing pipeline. Results from 3D classification are shown as central slices.

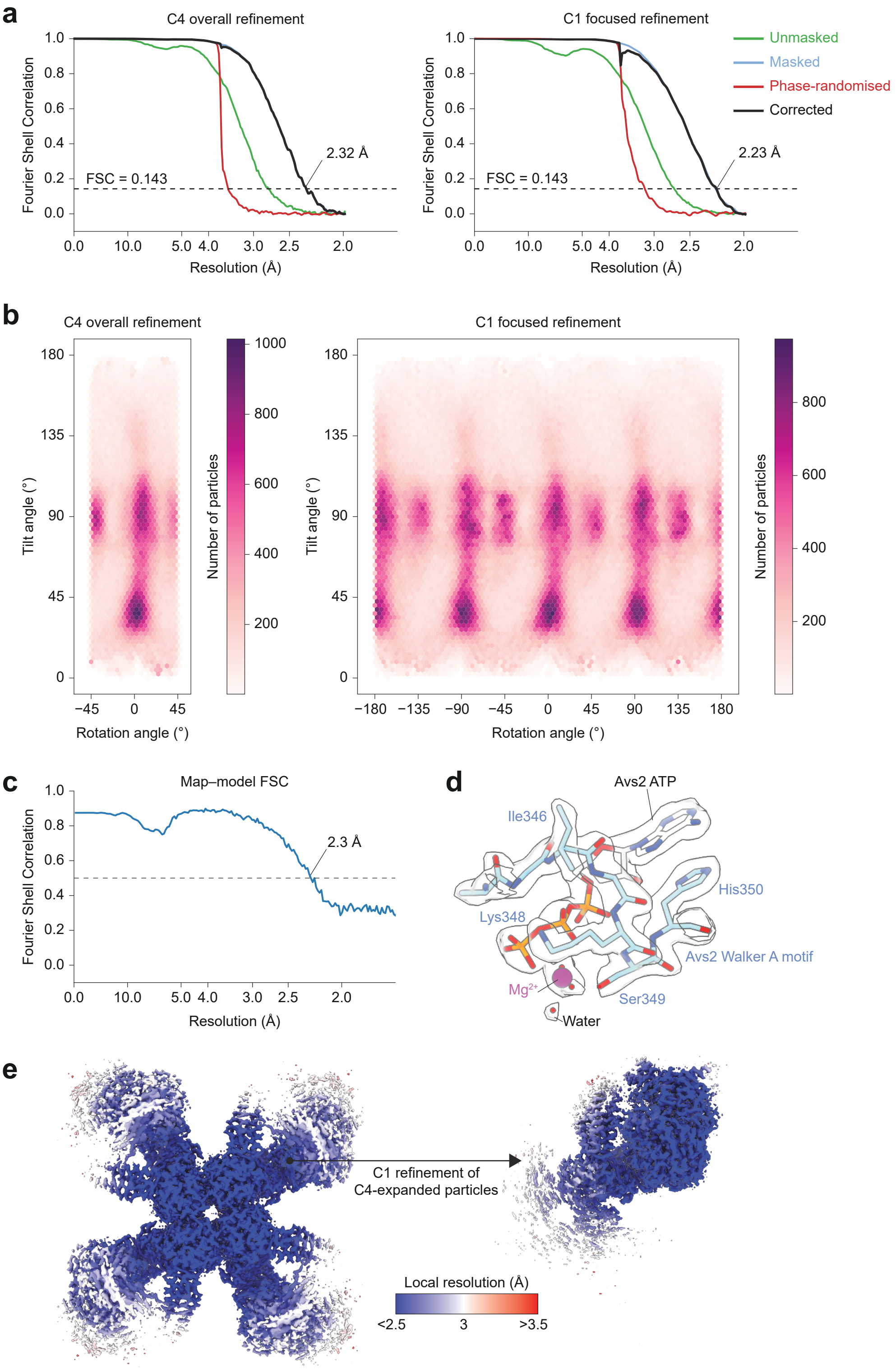


**Supplementary Fig. 4 | Cryo-EM data statistics and validation (related to Fig. 2).** **a**, Gold-standard Fourier-Shell Correlation (FSC) curves for the two final maps. **b**, Particle orientation distribution plots. Refinements performed with C4 symmetry assign rotation angles restricted to the range between –45 and +45 degrees. **c**, Map-to-model FSC curve calculated in PHENIX between the tetrameric model and a composite map. **d**, Cryo-EM density for the Walker A motif of EcAvs2 and the bound ATP molecule. **e**, Sharpened cryo-EM densities colored by local resolution as calculated using RELION.

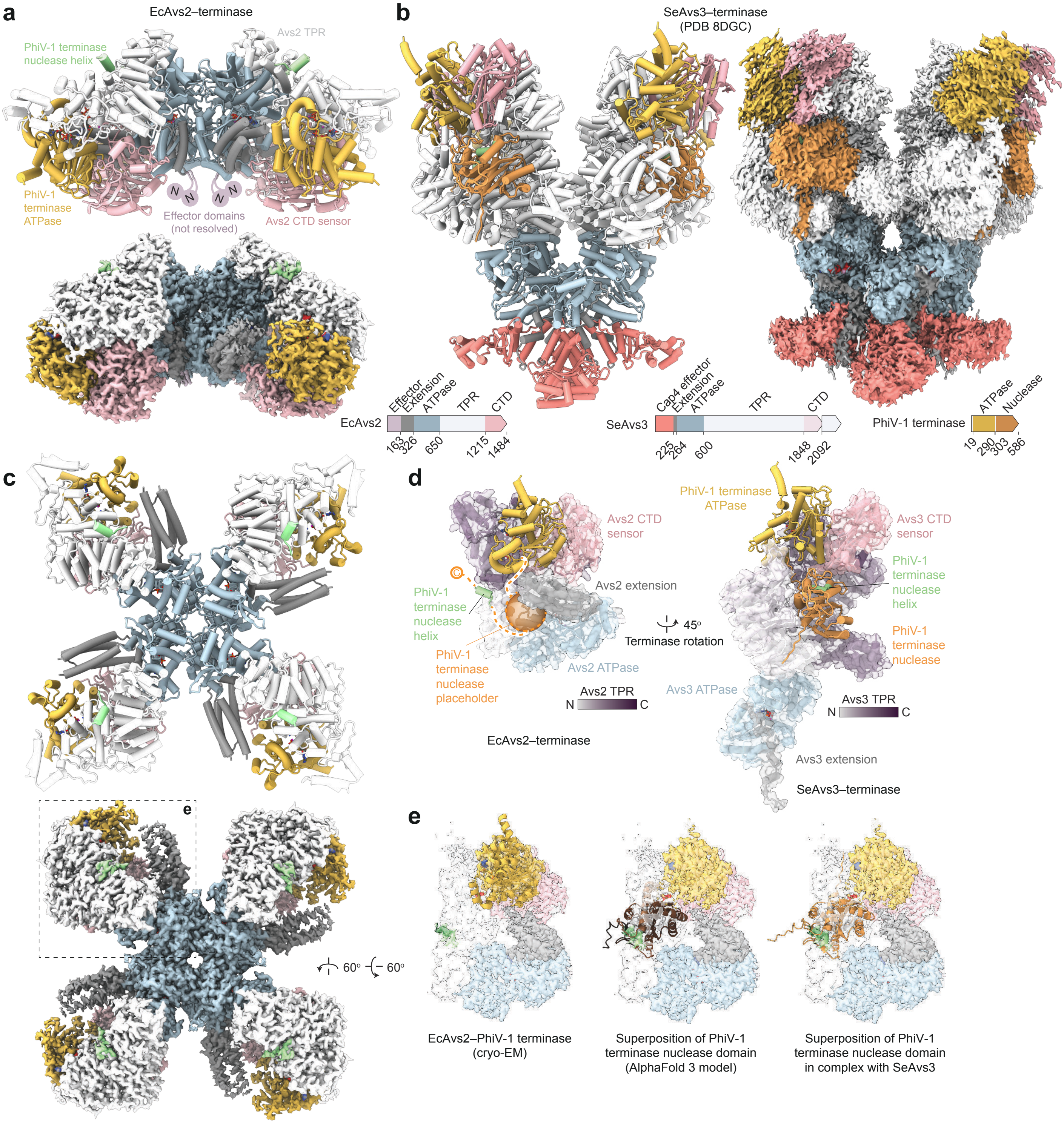


**Supplementary Fig. 5 | Comparison of terminase recognition by EcAvs2 and SeAvs3 (related to Fig. 2). a, b**, Side views of the atomic models and cryo-EM densities, colored by domain, for the EcAvs2–PhiV-1 (**a**) and SeAvs3–PhiV-1 (**b**) terminase complexes. **c**, Additional view of the EcAvs2–terminase complex atomic model and cryo-EM density. **d**, Structural comparison between the protomers of the EcAvs2–terminase complex and the SeAvs3–terminase complex (PDB 8DGC). The EcAvs2 TPR domain is colored from N- to C-terminus as indicated. The 11-amino acid nuclease helix is shown in green in both structures. In the EcAvs2 complex, the putative position of the remainder of the nuclease domain is indicated by an orange sphere. **e**, Structural superposition of the EcAvs2–terminase complex with an AlphaFold 3 model of the apo PhiV-1 terminase nuclease (middle) and the SeAvs3-bound terminase nuclease (right), aligned on the 11-residue nuclease helix.

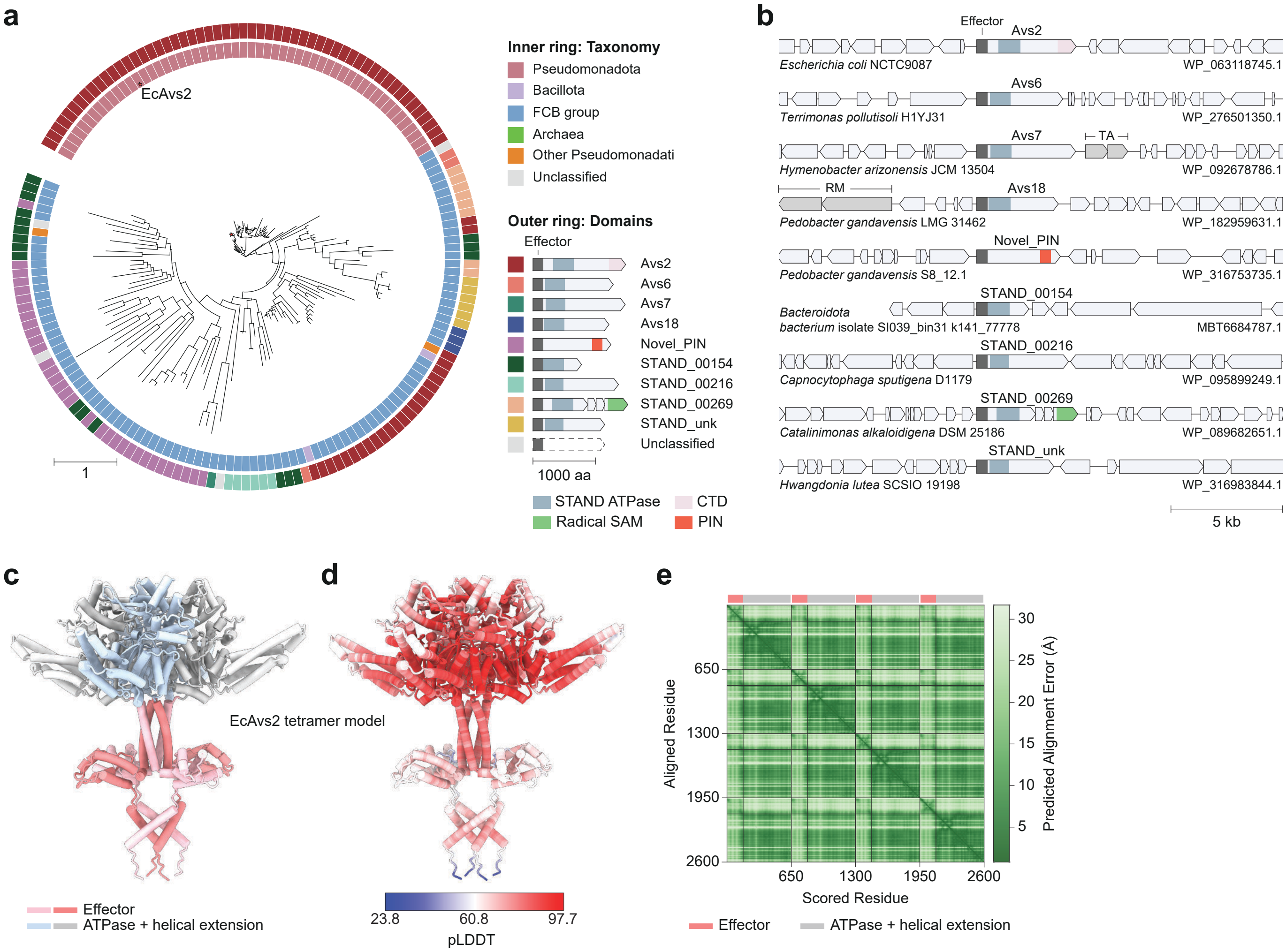


**Supplementary Fig. 6 | The EcAvs2 effector domain is present in diverse defense proteins. a**, Maximum likelihood tree of the EcAvs2 effector domain. **b**, Representative genomic neighborhood of 9 defense systems containing the effector. GenBank accessions correspond to the protein containing the effector. **c, d**, AlphaFold 3 models of an EcAvs2 effector–ATPase tetramer colored by domains (**c**) and pLDDT score (**d**). **e**, Predicted alignment error for the model in (c–d).

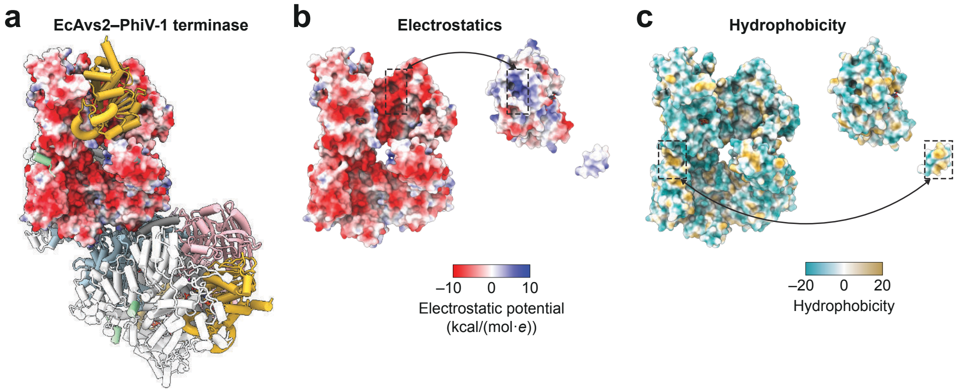


**Supplementary Fig. 7 | EcAvs2 recognizes phage terminases through shape and charge complementarity.** **a**, Tetrameric structure of EcAvs2, with one protomer rendered as a surface colored by electrostatic potential. **b, c**, Open-book views displaying the electrostatic surface potential (**b**) and hydrophobicity (**c**) of the interaction interface between EcAvs2 and the PhiV-1 terminase.

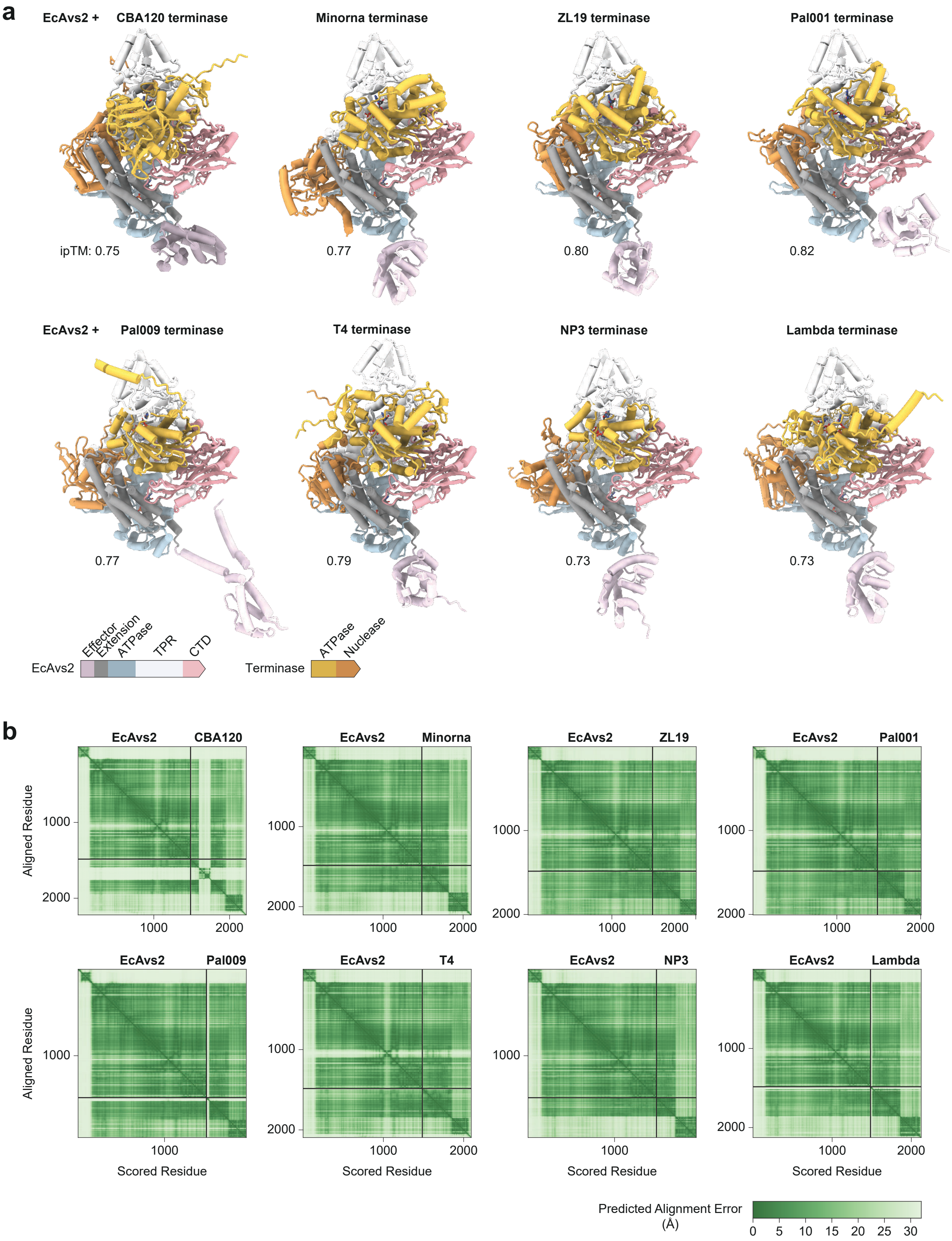


**Supplementary Fig. 8 | EcAvs2 is predicted to bind diverse terminases. a**, AlphaFold 3 models of EcAvs2 co-folded with eight phage terminases. ipTM, interface predicted template modeling score; values refer to the protein–protein interface and exclude ligands. **b**, Predicted alignment error of the models in (a).

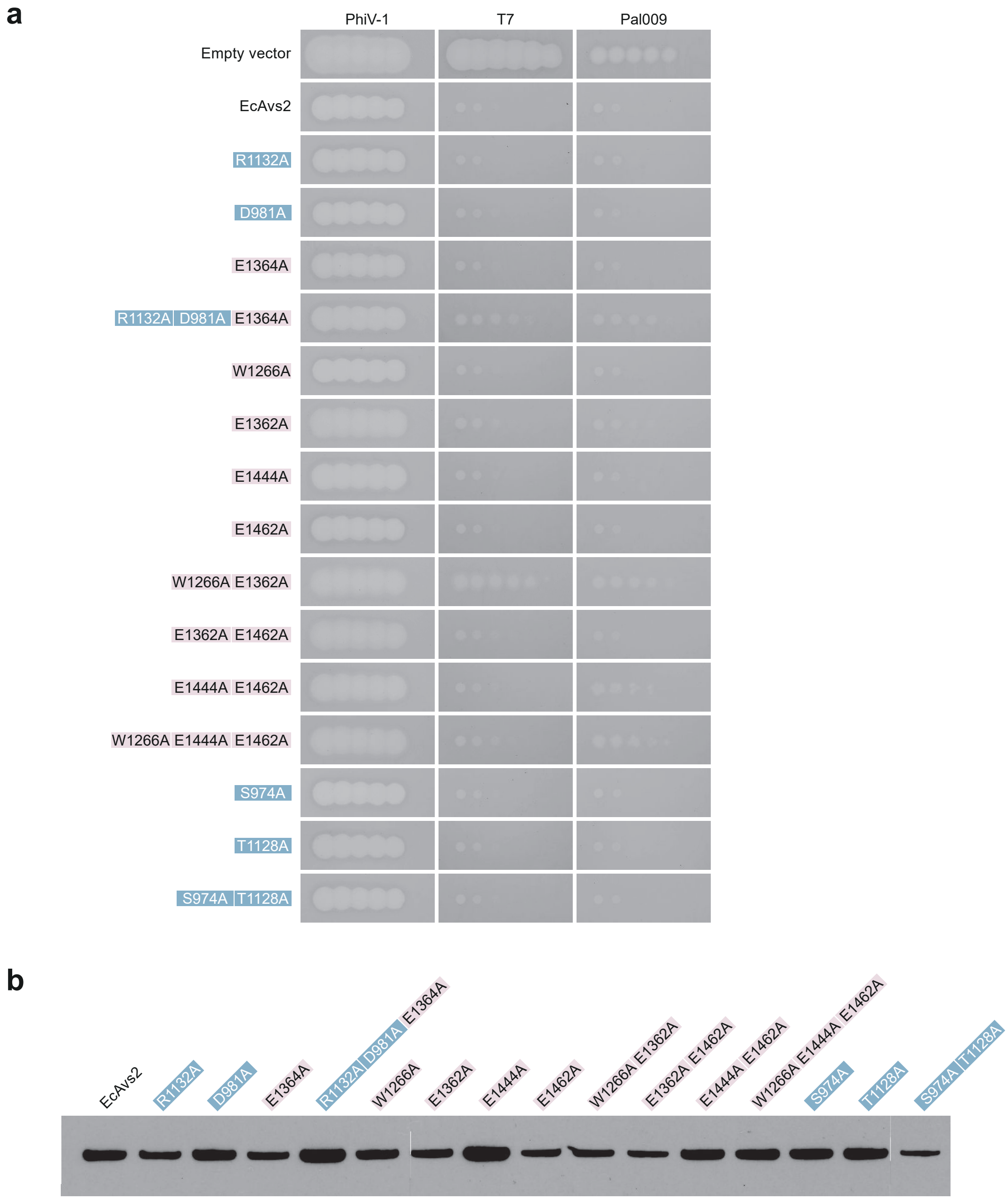


**Supplementary Fig. 9 | Validation of EcAvs2 mutants and protein expression (related to Fig. 3c).** **a**, Plaque assays of EcAvs2 interface mutants against phages PhiV-1, T7, and Pal009. **b**, Western blot of mutants in (a) heterologously expressed in *E. coli* with the native EcAvs2 promoter.

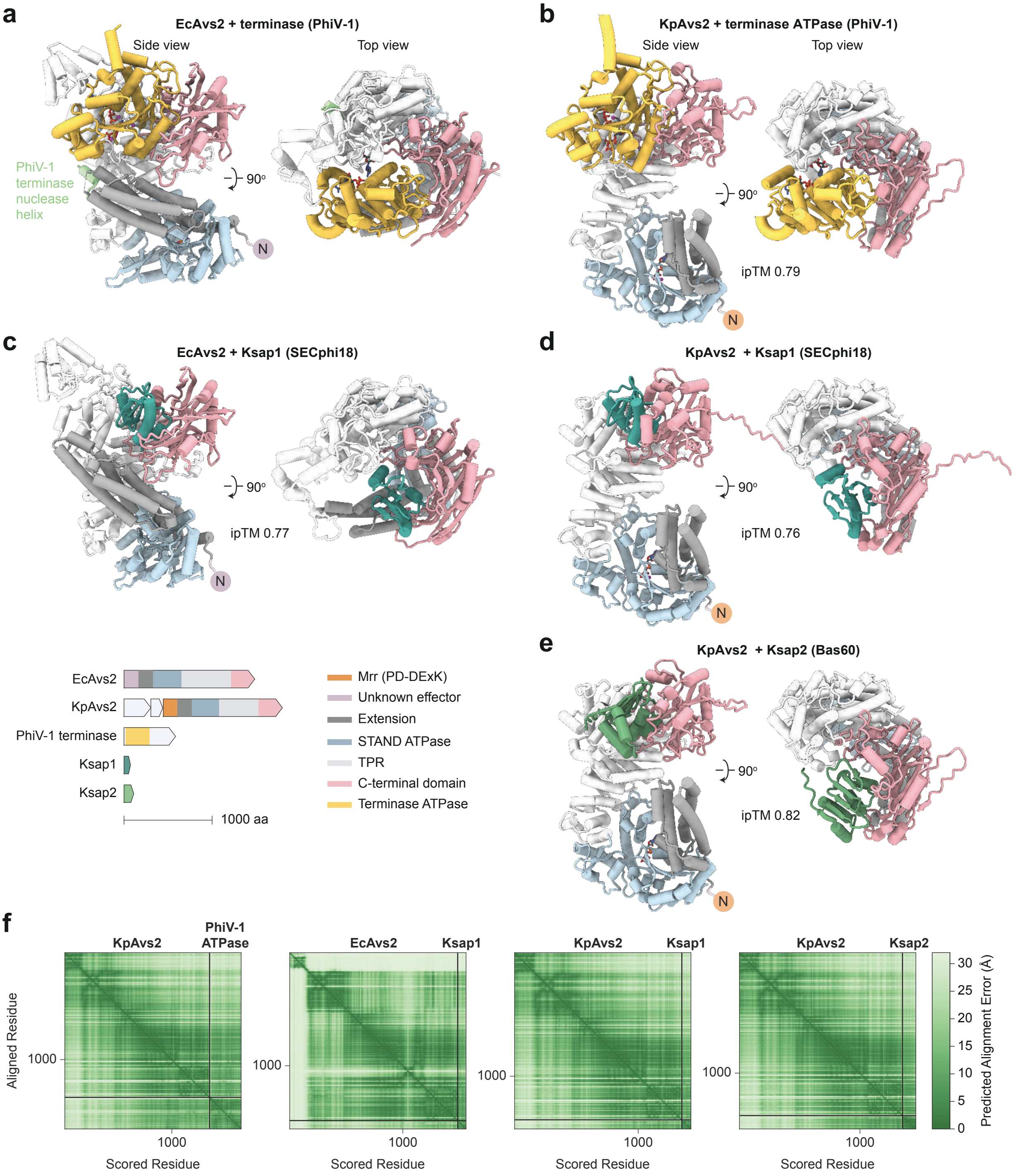


**Supplementary Fig. 10 | Comparison of terminase and Ksap1/2 binding to Avs2. a**, Cryo-EM structure of EcAvs2–PhiV-1 terminase. **b–e**, AlphaFold models of KpAvs2 and EcAvs2 bound to the terminase ATPase domain or Ksap1/2 activators. ipTM, interface predicted template modeling score; values refer to the protein–protein interface and exclude ligands. Predicted buried surface areas are as follows: EcAvs2-terminase (2833.5Å$^2$), KpAvs2-terminase (2508.9Å$^2$), EcAvs2-Ksap1 (1573.6Å$^2$), KpAvs2-Ksap1 (1481.6Å$^2$), and KpAvs2-Ksap2 (1637.2Å$^2$). **f**, Predicted alignment error for models in (b–e).

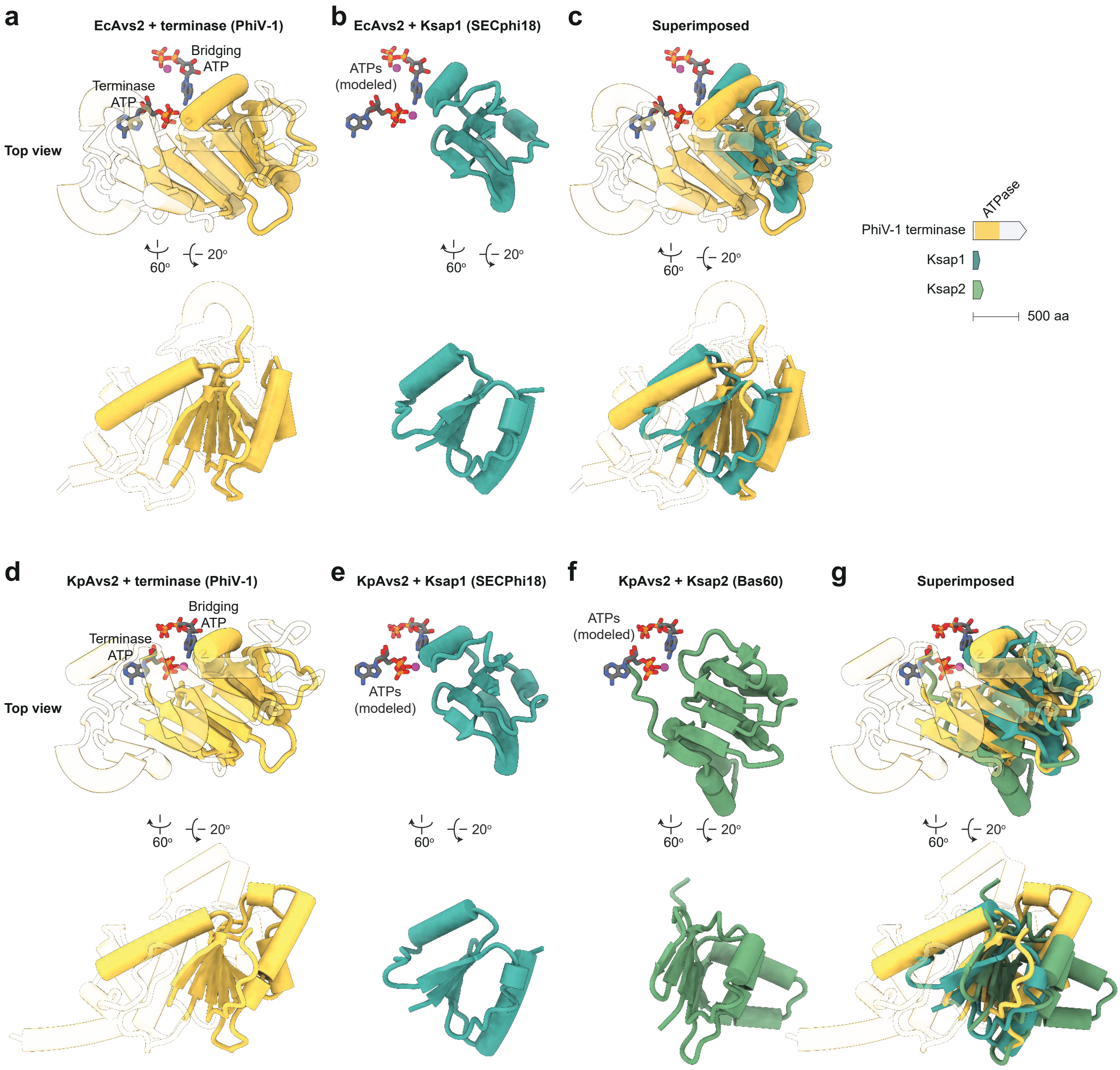


**Supplementary Fig. 11 | Structural comparisons of terminase and Ksap1/2. a**, Structure of the EcAvs2–PhiV-1 terminase complex. **b**, Model of the EcAvs2–Ksap1 complex, with ATP ligands from the PhiV-1 terminase structure shown for reference. **c**, Superposition of (a) and (b), aligned on EcAvs2. **d**, Model of the KpAvs2–PhiV-1 terminase complex. **e, f**, Models of KpAvs2–Ksap1 (**e**) and KpAvs2–Ksap2 (**f**), with ATP ligands from the PhiV-1 terminase model overlaid. **g**, Superposition of (d–f), aligned on KpAvs2.

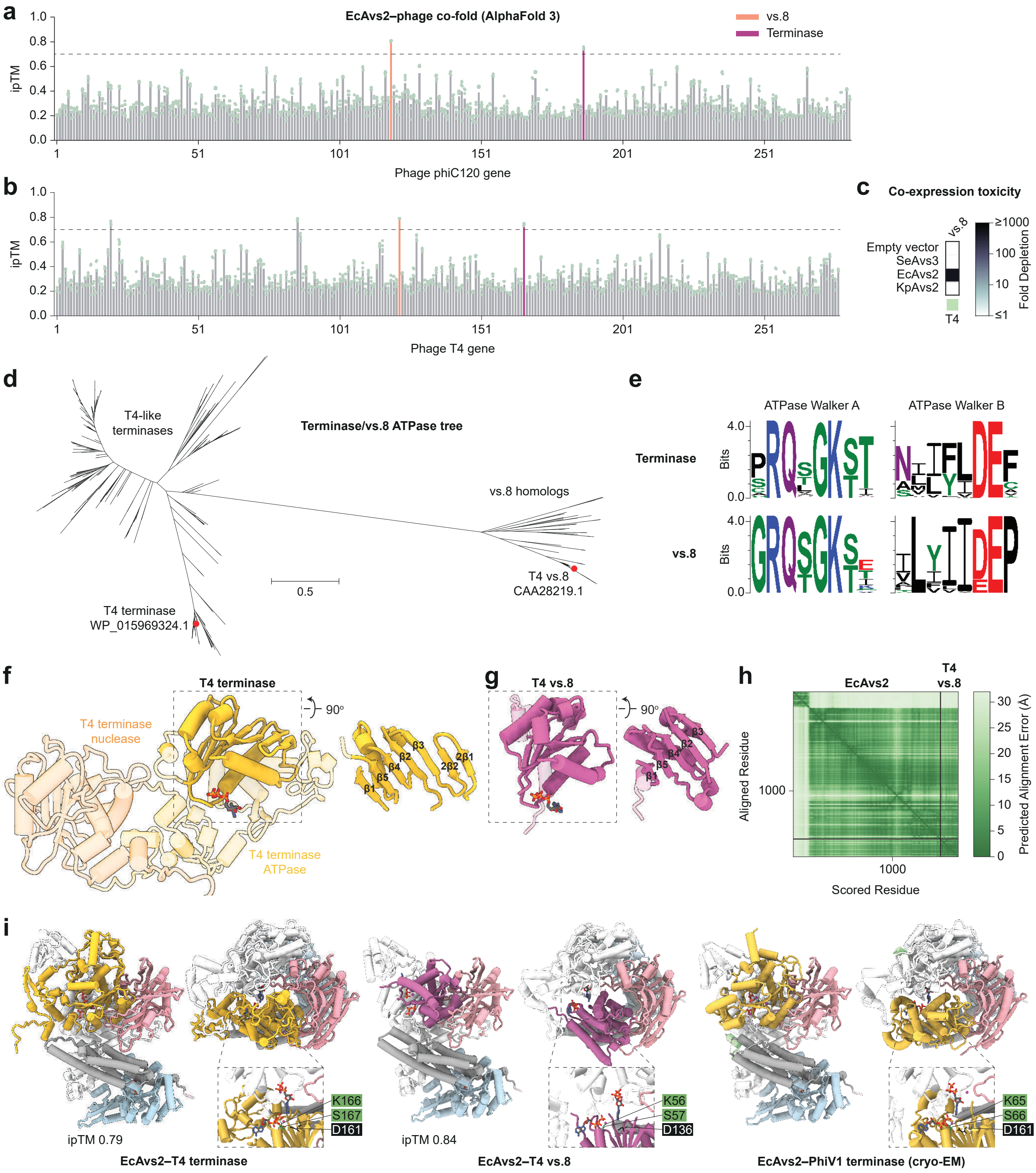


**Supplementary Fig. 12 | The ATPase-like protein vs.8 from phage T4 triggers EcAvs2-mediated toxicity. a, b**, AlphaFold 3 *in silico* screen for phage phiC120 (**a**) and T4 (**b**) proteins predicted to bind to EcAvs2. **c**, Toxicity of T4 vs.8 co-expressed with SeAvs3, EcAvs2, and KpAvs2. **d**, Maximum likelihood phylogenetic tree of vs.8 and the closest related phage terminase ATPase domains. **e**, Sequence logo of the ATPase Walker A and Walker B motifs of terminase and vs.8. **f, g**, AlphaFold 3 models of T4 terminase (**f**) and T4 vs.8 (**g**). **h**, Predicted alignment error for an AlphaFold 3 model of an EcAvs2–T4 vs.8 complex. **i**, Comparison of AlphaFold 3 models of EcAvs2–T4 terminase (left) and EcAvs2–T4 vs.8 (middle), together with the cryo-EM structure of EcAvs2–PhiV-1 terminase (right). Insets show terminase ATP and bridging ATP molecules, and green and black numbers denote conserved residues within the terminase ATPase Walker A and Walker B motifs, respectively. ipTM, interface predicted template modeling score; values refer to the protein–protein interface and exclude ligands.

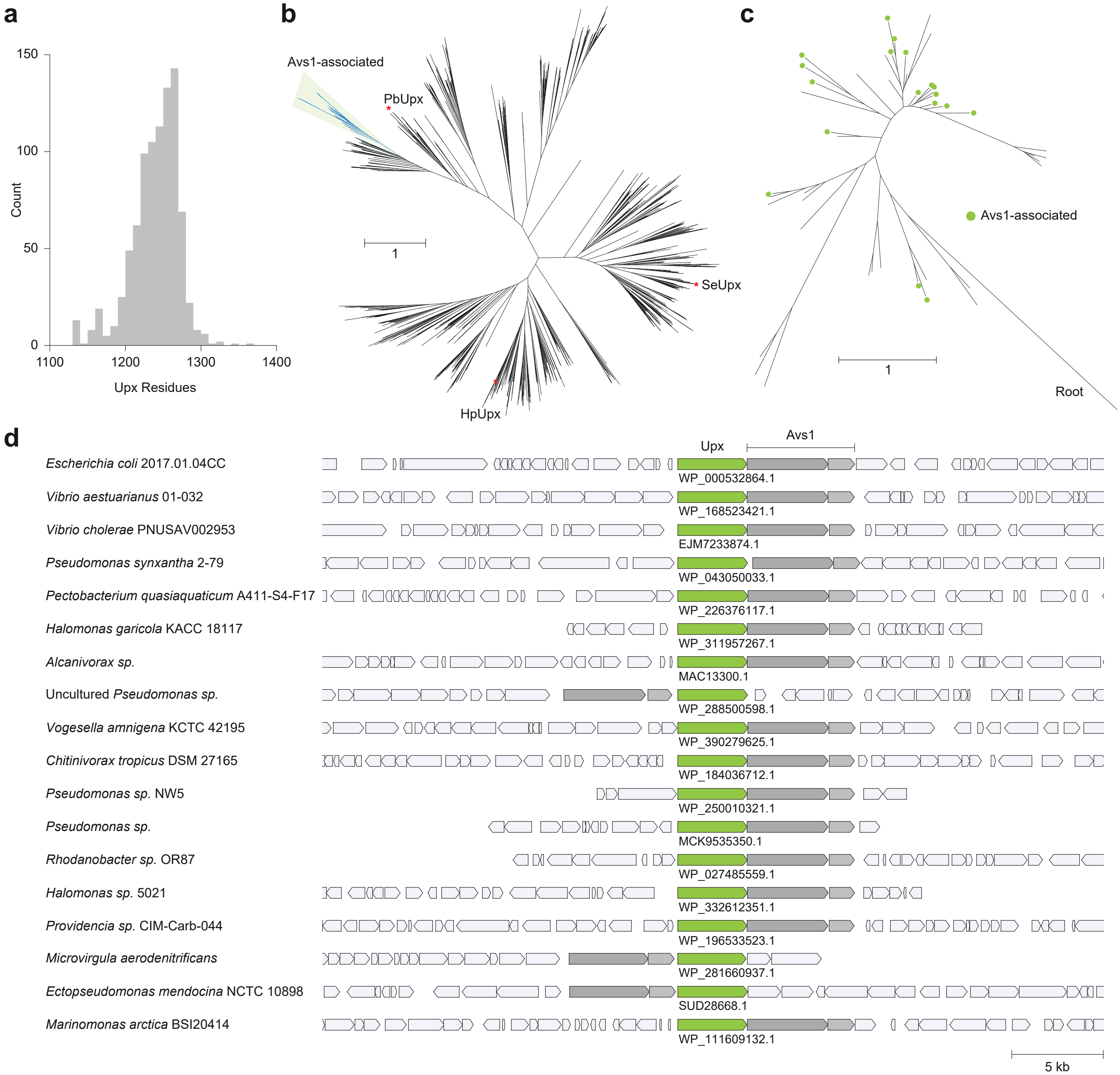


**Supplementary Fig. 13 | Computational analysis of Upx homologs (related to Fig. 4a). a**, Histogram of Upx protein length (median 1,245 amino acids). **b**, Maximum likelihood phylogenetic tree of Upx shown as an unrooted tree, with the Avs1-associated clade highlighted. **c**, Expanded view of the Avs1-associated clade. **d**, Genomic neighborhoods of 18 Avs1-associated Upx homologs.

**a** Terminase ATPase core β-sheet

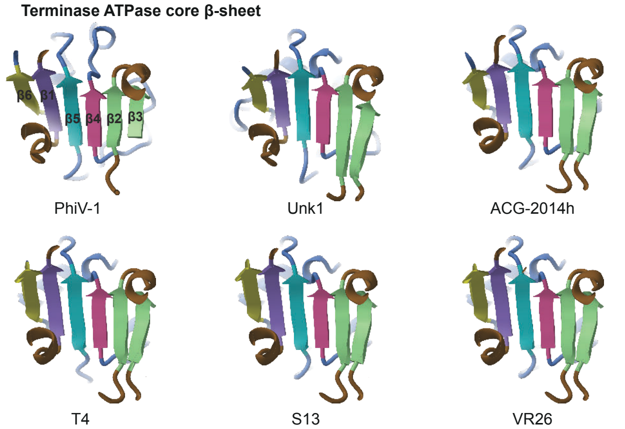


**b** Terminase ATPase core β-sheet color annotations

| Strand | PhiV-1 | Unk1 | ACG-2014h | T4 | S13 | VR26 |
|---|---|---|---|---|---|---|
| 1c | 54–59 | 133–137 | 75–79 | 155–160 | 165–170 | 155–160 |
| 2c | 85–89 | 162–168 | 105–110 | 185–190 | 195–200 | 185–190 |
| 3c | 141–145 | 210–215 | 153–158 | 233–238 | 243–248 | 233–238 |
| 4c | 157–161 | 229–233 | 171–175 | 251–255 | 261–265 | 251–255 |
| 5c | 196–201 | 261–266 | 203–208 | 281–286 | 291–296 | 281–286 |
| 6c | 220–224 | 293–296 | 231–234 | 309–313 | 319–323 | 309–313 |

**c**

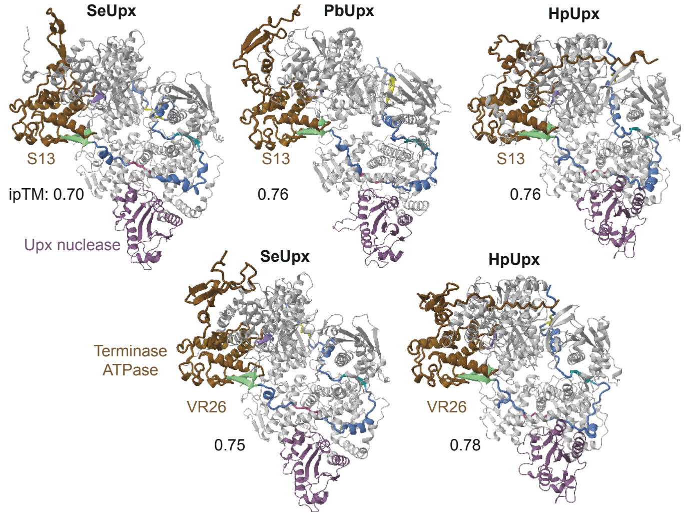


**d** Terminase (ACG-2014h)

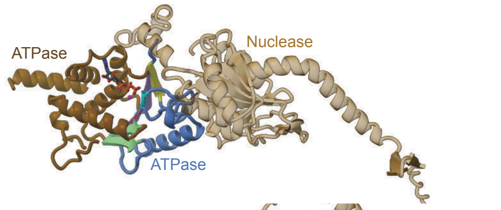


**e** Terminase (ACG-2014h) bound to PbUpx

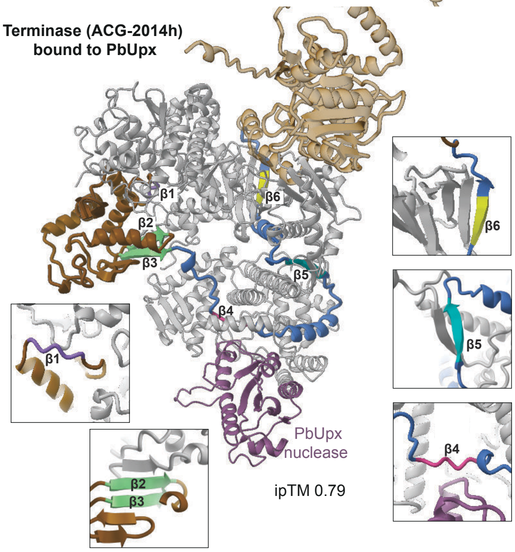


**Supplementary Fig. 14 | Structural models of the Upx–terminase binding interface (related to Fig. 4g–h). a**, Core β-sheet of six additional terminase ATPase domains, colored according to the scheme in Fig. 4g. **b**, Residue numbers of the β-strands in (a). **c**, Five additional AlphaFold 3 models of SeUpx, HpUpx, or PbUpx in complex with a terminase ATPase domain. **d, e**, Comparison of the AlphaFold 3 structures of the phage ACG-2014h terminase in the apo state (**d**) and in complex with PbUpx (**e**). ipTM, interface predicted template modeling score; values refer to the protein–protein interface and exclude ligands.

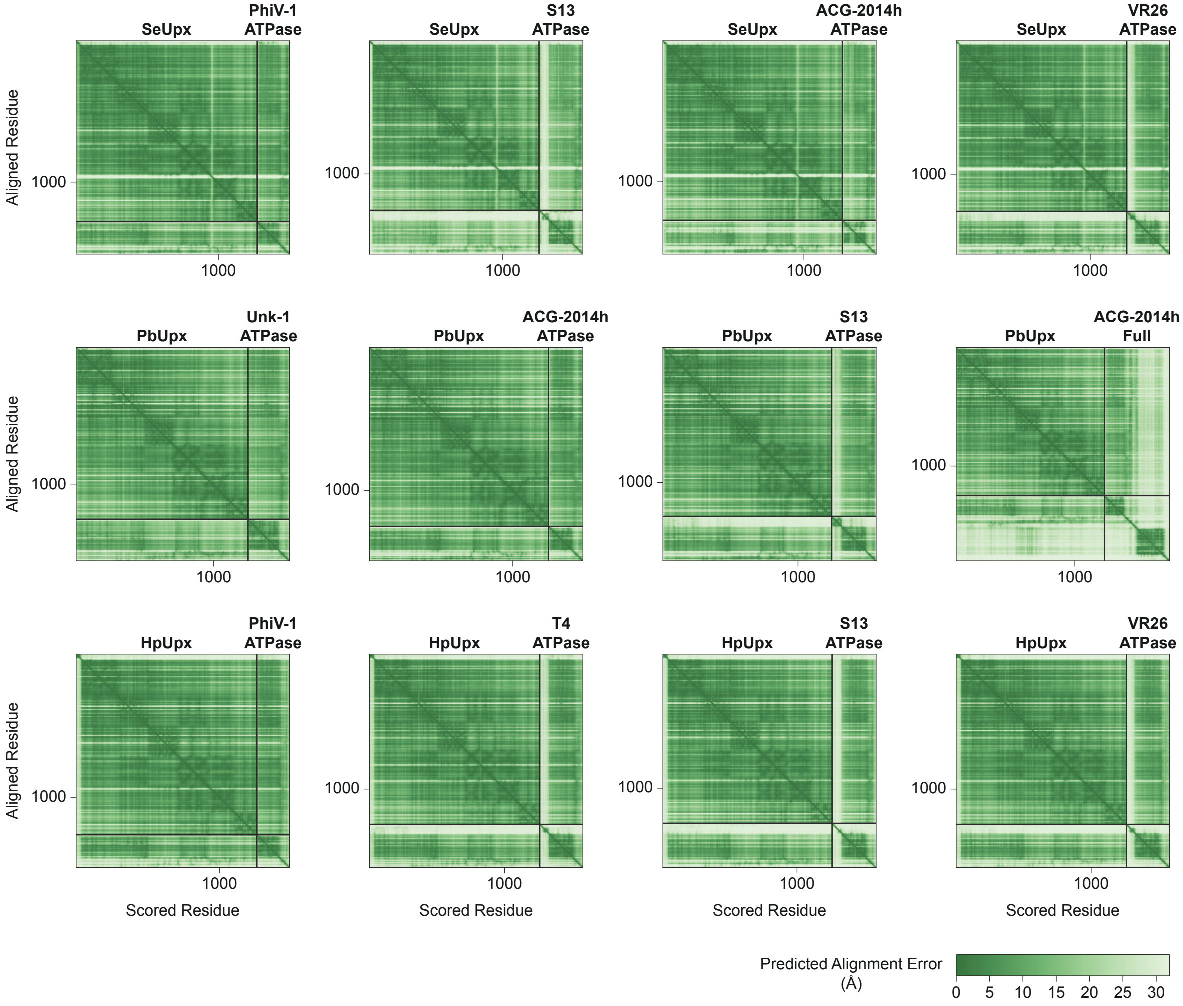


**Supplementary Fig. 15 | Predicted alignment error of the AlphaFold 3 models of Upx–terminase complexes shown in Supplementary Fig. 14.**
